# Rhinovirus C replication is associated with the endoplasmic reticulum and triggers cytopathic effects in an *in vitro* model of human airway epithelium

**DOI:** 10.1101/2021.04.16.440244

**Authors:** Talita B. Gagliardi, Monty E. Goldstein, Daniel Song, Kelsey M. Gray, Jae W. Jung, Kimberly M. Stroka, Gregg A. Duncan, Margaret A. Scull

## Abstract

The clinical impact of rhinovirus C (RV-C) is well-documented; yet the viral life cycle remains poorly defined. Thus, we characterized RV-C15 replication at the single-cell level and its impact on the human airway epithelium (HAE) using a physiologically-relevant *in vitro* model. RV-C15 replication was restricted to ciliated cells where viral RNA levels peaked at 12 hours post-infection (hpi), correlating with elevated titers in the apical compartment at 24 hpi. Notably, infection was associated with a loss of polarized expression of the RV-C receptor, cadherin-related family member 3. Visualization of double-stranded RNA (dsRNA) during RV-C15 replication revealed two distinct replication complex arrangements within the cell, likely corresponding to different time points in infection and correlating with the formation of large intracellular vesicles. To further define RV-C15 replication sites, we analyzed the expression of giantin, phosphatidylinositol-4- phosphate, and calnexin, as well as the colocalization of these markers with dsRNA. Fluorescence levels of all three cellular markers were elevated during infection and altered giantin distribution further indicated Golgi fragmentation. However, unlike previously characterized RVs, the high ratio of calnexin-dsRNA colocalization implicated the endoplasmic reticulum as the primary site for RV-C15 replication in HAE. RV-C15 infection was also associated with elevated stimulator of interferon genes (STING) expression, facilitating replication, and the induction of incomplete autophagy, a mechanism used by other RVs to promote non-lytic release of progeny virions. Finally, RV-C15 infection resulted in a temporary loss in epithelial barrier integrity and the translocation of tight junction proteins while a reduction in mucociliary clearance indicated cytopathic effects on epithelial function. Together, our findings identify both shared and unique features of RV-C replication compared to related rhinoviruses and define the impact of RV-C on both epithelial cell organization and tissue functionality – aspects of infection that may contribute to pathogenesis *in vivo*.

**Author summary:** Rhinovirus C has a global distribution and significant clinical impact – especially in those with underlying lung disease. Although RV-C is genetically, structurally, and biologically distinct from RV-A and -B viruses, our understanding of the RV-C life cycle has been largely inferred from these and other related viruses. Here, we performed a detailed analysis of RV-C15 replication in a physiologically-relevant model of human airway epithelium. Our single-cell, microscopy-based approach revealed that – unlike other RVs – the endoplasmic reticulum is the primary site for RV- C15 replication. RV-C15 replication also stimulated STING expression, which was proviral, and triggered dramatic changes in cellular organization, including altered virus receptor distribution, fragmented Golgi stacks, and the induction of incomplete autophagy. Additionally, we observed a loss of epithelial barrier function and a decrease in mucociliary clearance, a major defense mechanism in the lung, during RV-C15 infection. Together, these data reveal novel insight into RV-C15 replication dynamics and resulting cytopathic effects in the primary target cells for infection, thereby furthering our understanding of the pathogenesis of RV-C. Our work highlights similar, as well as unique, aspects of RV-C15 replication compared to related pathogens, which will help guide future studies on the molecular mechanisms of RV-C infection.

## Introduction

Rhinoviruses (RVs) are responsible for over 40% of respiratory virus infections in the human population [1–4]. Although well known as etiologic agents of the common cold, rhinoviruses can also infect the lower respiratory tract causing bronchiolitis or pneumonia and are a leading cause of virus-induced exacerbations in acute and chronic lung disease [5–8]. No vaccine or direct-acting antiviral is currently available due in part to the diversity of RVs in circulation, with over 160 genotypes identified [9–10]. These genotypes comprise three species (RV-A, RV-B, and RV-C) where RV-A and RV-C are the most prevalent and RV-C is associated with more severe clinical disease, especially in children [4, 11, 12]. Indeed, RV infection during the first year of life has been associated with wheezing episodes and is considered a risk factor for the development of asthma [8, 13, 14].

RV-C was discovered in 2006 [15] and compared to previously defined rhinovirus species, is unique at the genetic [16], structural [17], and biological level [18]. While all RVs (-A, -B, and -C) infect airway epithelial cells, RV-C uses a different host protein, cadherin-related family member 3 (CDHR3), to mediate particle uptake [19]. The restricted expression of CDHR3 to ciliated cells in the upper and lower airway epithelium [20–23] limits the cellular tropism of RV-C, compared to other RVs that utilize low-density lipoprotein receptor (LDLR) or intercellular adhesion molecule (ICAM)-1 as receptors [18]. Notably, a non-synonymous single nucleotide polymorphism (SNP; rs6967330[A]) that yields stabilized CDHR3 protein expression at the cell surface is a causal variant for early childhood asthma with severe exacerbations [20]. Subsequent investigation has associated the CDHR3 asthma risk allele with heightened risk of respiratory tract illness with RV-C, but not other viruses [21]. More recently, stimulator of interferon genes (STING), a key adapter protein for cytosolic DNA-sensing pathways, was found to play a proviral role in RV-A and -C, but not -B, replication [24]. Thus, mechanisms of infection and replication are not always conserved between rhinovirus species.

Despite these advances, details of the RV-C life cycle and underlying mechanisms that contribute to pathogenesis remain scarce. While such studies remain hampered by the absence of immortalized cell lines or an *in vivo* mouse model that is naturally susceptible and highly permissive for RV-C replication, previous reports demonstrate that *ex vivo* tissue and primary airway cultures support infection [22, 25]. Nonetheless, aside from receptor usage and cellular tropism, little is known about RV-C interactions with primary airway epithelial cells, the principal target for infection. Here we utilized both single-cell, microscopy-based analyses, and culture-wide measurements, to investigate the details of RV-C15 replication in an *in vitro* model of human airway epithelium (HAE). Our data identify the endoplasmic reticulum (ER) as the primary site for RV-C15 replication and demonstrate the impact of infection on the structural integrity of ciliated cells and epithelial barrier function, thereby identifying both unique and shared features of RV-C amongst related viruses.

## Results

### 1. RV-C15 replicates in ciliated cells, yielding changes in CDHR3 expression

Pseudostratified models of HAE at air-liquid interface are permissive for RV-C replication [25, 26]. To detail the kinetics of RV-C15 replication in this model at higher-resolution, we inoculated HAE at 34°C with 10^10^ RV-C15 RNA copies and quantified viral RNA intracellularly as well as in both the apical and basolateral compartments over time (**Fig 1A and 1B**). The dynamics of RV-C15 replication were similar in HAE from two different donors, where cell-associated RV- C15 RNA levels increased during the first 12 hours post-infection (hpi). This correlated with the detection of double-stranded (ds) RNA, a marker of viral replication, by immunofluorescence (IF; **S1A Fig**) and was in line with peak viral release into the apical chamber at 24 hpi (**Fig 1A and 1B**). The lack of RV-C15 RNA in basolateral supernatants confirmed polarized release of RV-C15 to the airway lumen (**Fig 1A and 1B**), consistent with a previous report in nasal cells [27] and the clinical manifestation of RV-C-mediated disease.

**Figure 1.**
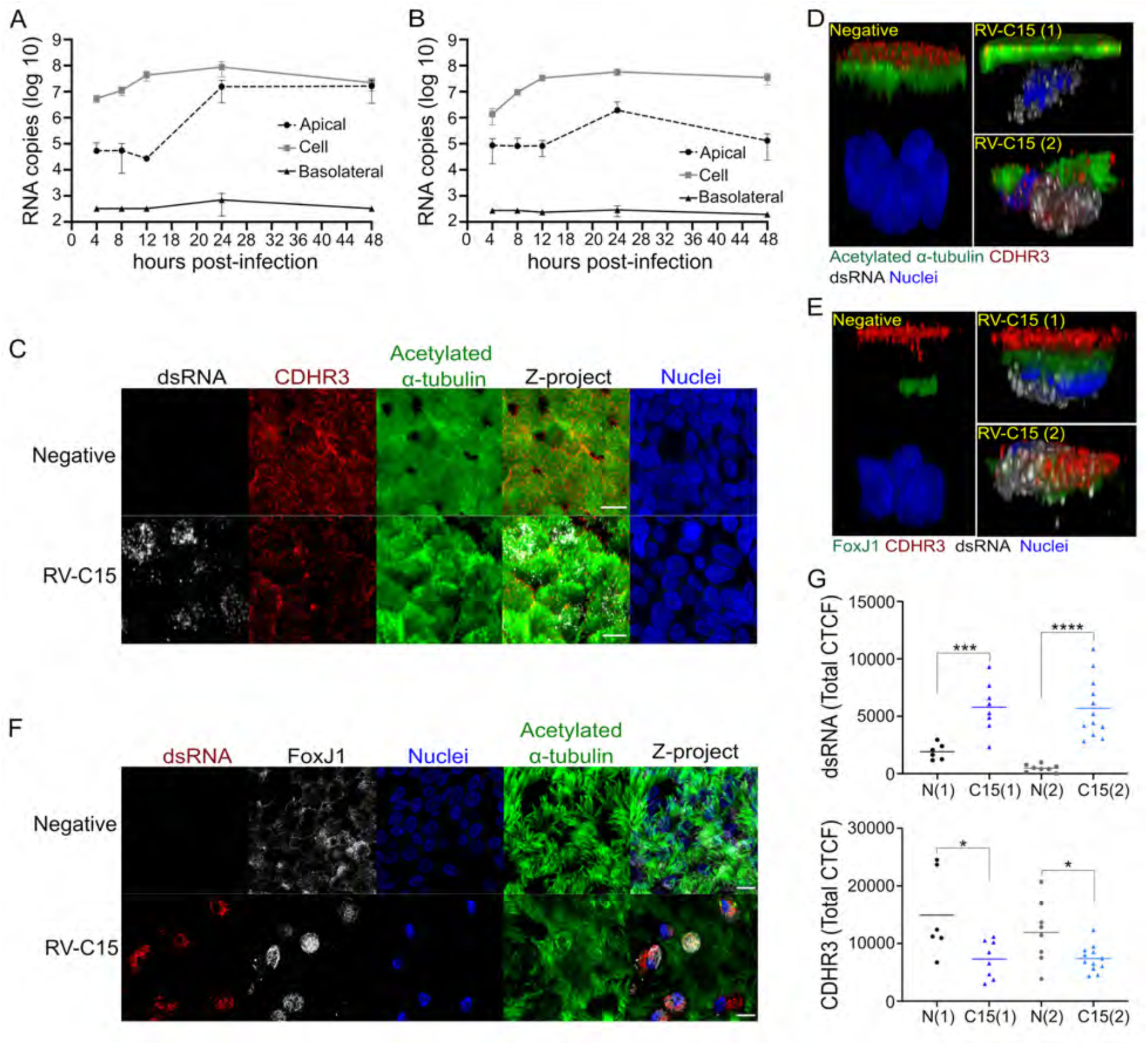
RV-C15 replicates in ciliated epithelial cells, leading to decreased CDHR3 levels. **A and B:** Multi-cycle RV-C15 growth delineated at 34°C in HAE (**A:** donor 1; **B:** donor 2). **C-E:** Immunofluorescence detection of CDHR3 (red) in non-infected or RV-C15-infected (dsRNA+; gray) ciliated cells identified by either acetylated α-tubulin (green; **C and D**) or FoxJ1 (green; **E**) at 12 hpi. **C:** scale bar = 10µm. **D and E:** 3D visualization; z-stacks were at 1µm of thickness. **F:** Immunofluorescence detection of RV-C15 replication (dsRNA; red) in ciliated cells (FoxJ1, gray) with/or without motile cilia (acetylated α-tubulin, green) at 12 hpi (scale bar = 10µm). **G:** Quantification of dsRNA and CDHR3 fluorescence levels (CTCF) in non-infected (N) and RV- C15-infected (C15; dsRNA+) ciliated cells following immunofluorescence staining at 12 hpi. Graphs show dsRNA and CDHR3 total CTCF (line = mean) quantified in HAE using FoxJ1 (N(1) and C15(1)) or acetylated α-tubulin (C15(2) and N(2)) as a marker of ciliated cells. Statistical analysis was done using the Mann-Whitney U test (Two-tailed; 0.95% confidence interval; *p<0.05, ***p<0.001, ****p<0.0001).

Given that HAE cultures were permissive for RV-C15 replication, we next sought to investigate cell tropism in our system. Visualization of the RV-C receptor, CDHR3, in non-infected HAE by IF revealed expression at the apical surface in cells that were also positive for Forkhead box protein J1 (FoxJ1; known to promote ciliogenesis) or acetylated alpha-tubulin (a marker of mature ciliated cells; **Fig 1C, 1D, and 1E**). Notably, the corrected total cellular fluorescence (CTCF) levels of CDHR3 had a limited relationship with FoxJ1 but were strongly correlated with acetylated alpha-tubulin levels, in line with increased expression of CDHR3 during differentiation (**S1B and S1C Fig)** [21]. Corroborating these data and previous research reporting ciliated cell tropism for RV-C [22, 25], we detected dsRNA primarily in FoxJ1(+) cells, with some cells also staining positive for acetylated alpha-tubulin (**Fig 1F and S1D Fig**). However, the visibility of the cilia in these cells was diminished (**Fig 1F**) and the 3D view suggested CDHR3 was internalized in some dsRNA(+) cells, together with a more diffuse distribution of acetylated alpha-tubulin and FoxJ1 (**Fig 1D and 1E**).

To better understand the dynamics of FoxJ1, acetylated alpha-tubulin, and CDHR3 during infection, we analyzed global protein expression by Western blot (WB) and quantified fluorescence levels following immunostaining specifically in infected cells. Beyond the intrinsic variation expected across cultures in this model system, neither FoxJ1 nor acetylated alpha-tubulin protein levels were dramatically altered over the course of infection while CDHR3 levels fluctuated (**S1E, S1F Fig**). Subsequent single-cell analysis of acetylated alpha-tubulin and FoxJ1 fluorescence levels corroborated our WB data (**S1G, S1H Fig**); however, in contrast to the global elevation of CDHR3 protein in RV-C15-infected HAE at 12 hpi, CDHR3 fluorescence levels decreased significantly in cells with active viral replication (**Fig 1G**).

### 2. RV replication complex distribution is associated with vesicle formation

Picornaviruses, including RV-A and -B, replicate in association with cellular membranes, leading to the formation of double-membrane vesicles known as replicative complexes [28–31]. Thus, we investigated the distribution of replication complexes in RV-C15-infected HAE cultures by visualization of viral dsRNA and the impact of infection on intracellular membrane organization by transmission electron microscopy (TEM). While non-infected HAE were negative for dsRNA, as expected (**Fig 2A**), dsRNA in RV-C15-infected cultures was found in either a perinuclear (**Fig 2B**) or “ring-like” disposition closer to apical and basolateral membranes (**Fig 2C**). Neither phenotype was specific for RV-C15, however, as HAE infected with either RV-A16 or RV-A2 revealed similar dsRNA profiles (**S2A-D Fig**). Notably, the nuclei in cells with the “ring-like” dsRNA pattern were found near the apical cell surface instead of the usual basolateral location in polarized columnar cells (**Fig 2C, S2B, S2D Fig**), possibly a result of large intracellular vesicle formation (**S2E and S2F Fig**).

**Figure 2.**
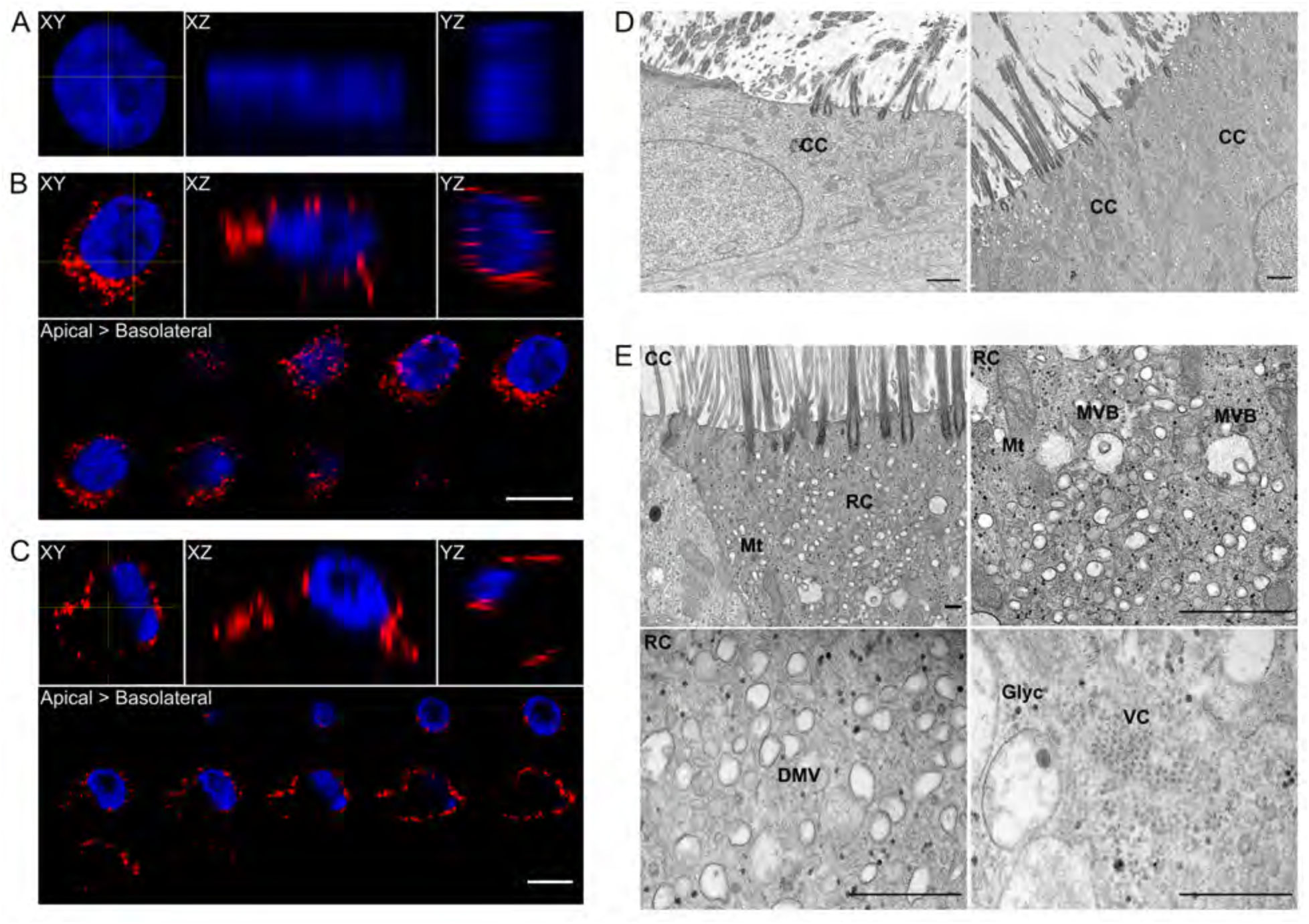
Detection of RV-C15 replication complexes in HAE. **A-C:** Orthogonal views (XY, XZ and YZ planes; yellow lines show the location of XZ and YZ views on the XY plane) from non- infected **(A)** and RV-C15-infected (**B-C**) cells immunostained for dsRNA (red) and nuclei (blue) at 12 hpi (z-stacks at 1µm of thickness; scale bar = 10µm). **B-C**: Z-stacks (1µm of thickness) from RV-C15-infected HAE show dsRNA (red) detection by immunofluorescence with perinuclear **(B**) or “ring-like” disposition close to the plasma membrane (**C**). **D-E:** TEM of non-infected (**D**) and RV-C15-infected (**E)** HAE fixed at 12 hpi (scale bar = 5µm). **D:** Visualization of ciliated cells *(CC*) in non-infected HAE (**D –** *left panel)* occasionally revealed small vesicles spread through the cytoplasm (**D** *– right panel*). **E:** Larger vesicles were detected in ciliated cells (*CC*) from RV-C15- infected HAE that resembled ’replicative complexes’ (*RC*; **E** *– upper-left panel*) observed during RV-A and RV-B infection [30]. These vesicles were detected in close proximity to mitochondria (*Mt*) and multi-vesicular bodies (*MVB*; **E** *– upper-right panel*), and were found to have double- membranes (DMV; **E** *– lower-left panel*). Clusters of electron-dense structures in the cytoplasm of RV-C15-infected HAE were seen, reminiscent of viral crystals (VC; **E** *– lower-right panel*) described for other picornaviruses [32, 33]. Glyc = glycogen.

Subsequent TEM analysis in non-infected HAE revealed a differentiated epithelium with no overt cytopathic effects (**Fig 2D – left panel**) and a limited number of vacuoles in the cytoplasm (**Fig 2D- right panel**). In contrast, in RV-C15-infected HAE, we identified ciliated cells with many small vesicles clustered near multivesicular bodies, mitochondria, and electron-dense structures similar to β-particles of glycogen (**Fig 2E – upper-left and upper-right panels, respectively**). Under higher magnification, we were able to confirm these small vesicles had a double-membrane (**Fig 2E – lower-left panel**) similar to replicative complexes previously described for RV-A and RV-B [30]. Interestingly, some ciliated cells in RV-C15-infected HAE also contained electron-dense structures in their perinuclear region similar to the “viral crystals” observed in cells infected by other picornaviruses (**Fig 2E – lower-right panel**) [32, 33].

### 3. RV-C15 infection triggers fragmentation of Golgi stacks and induces PI4P

Different cellular membranes can contribute to the formation of replication organelles during viral infection; however, the Golgi is the main source reported for many picornaviruses, including rhinoviruses [31, 34]. Consequently, viral replication is associated with fragmentation of the Golgi stacks and changes in expression of phosphatidylinositol-4-phosphate (PI4P), a Golgi resident lipid [31, 34, 35]. To determine if RV-C replication induced similar effects, we inoculated HAE with RV-C15 – or RV-A16 and RV-A2 as positive controls [35, 36] – and analyzed the Golgi by detection of giantin expression at 12 hpi. While giantin visualization revealed a compact structure close to the nucleus in non-infected HAE (**Fig 3A**), giantin signal was spread throughout the cytoplasm in cells with evidence of active viral replication for all RVs tested (**Fig 3A**). The fragmentation of Golgi structures inferred from our IF data was further supported by TEM, where, compared to the non-infected control, Golgi stacks were barely visible in cells from RV-C15- infected HAE cultures (**Fig 3B**).

**Figure 3.**
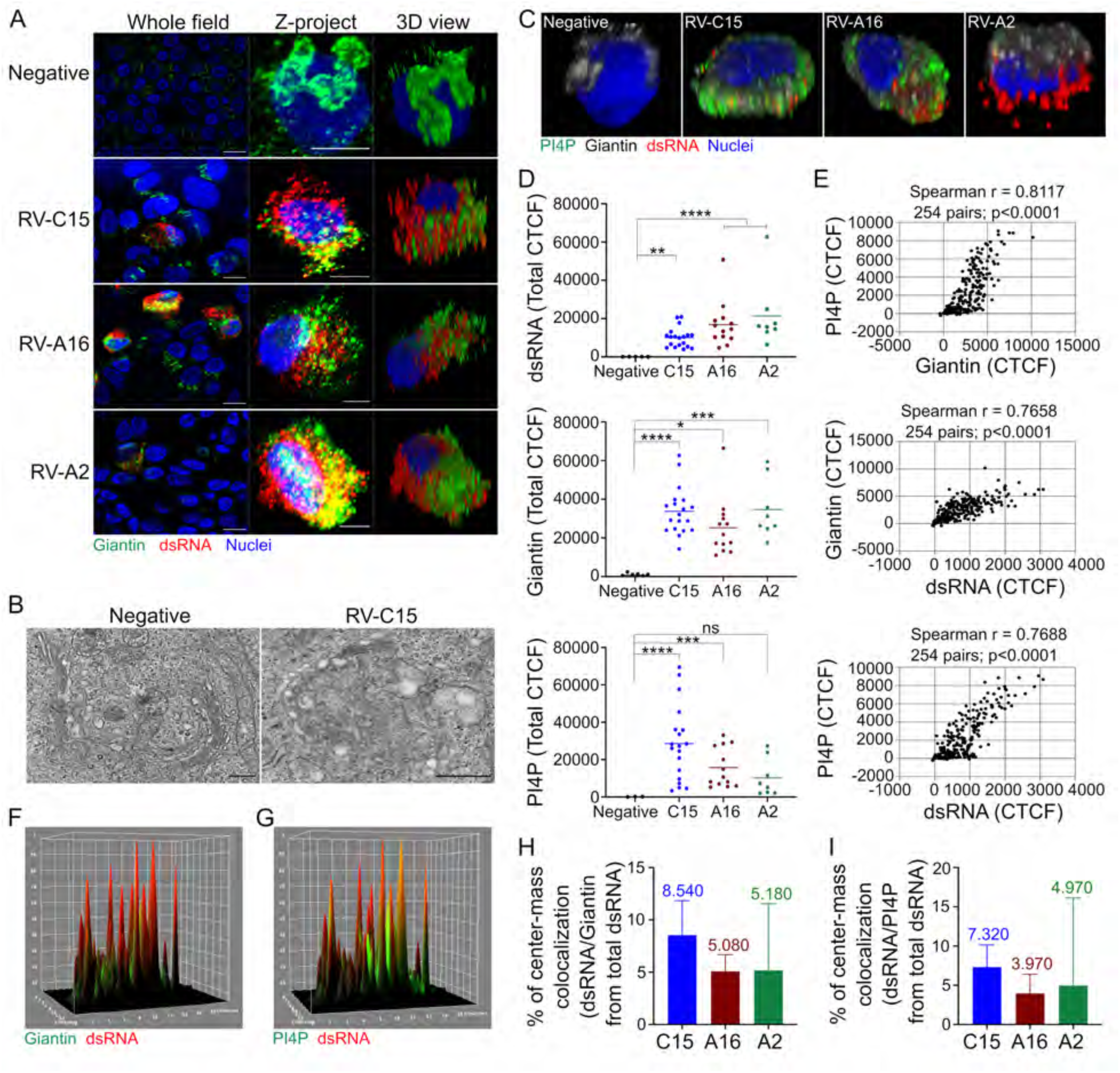
Neither the Golgi nor PI4P-positive vesicles are the main site for RV-C15 replication in HAE. **A**: Visualization of giantin (Golgi marker, green; nuclei, blue) in non-infected HAE and in HAE infected with RV-C15, RV-A16, or RV-A2 (dsRNA+, red) by immunofluorescence (z-stacks at 1µm of thickness; scale bar = 10µm) at 12hpi. **B:** Golgi stacks observed by TEM in ciliated cells from non-infected (*left panel*) and RV-C15-infected HAE (*right panel*) at 12 hpi (scale bar = 2µm). **C**: 3D view of PI4P (green) and giantin (gray) in non-infected or rhinovirus-infected (dsRNA+, red) HAE (nuclei, blue) detected by immunofluorescence at 12hpi (z-stacks at 1µm of thickness). **D:** Quantification of fluorescence levels (CTCF; line = mean) of dsRNA, giantin, and PI4P in non-infected HAE and HAE infected with RV-C15, RV-A16, or RV-A2 (dsRNA+ cells) at 12 hpi. Statistical analysis was done using the Kruskal-Wallis test following Dunn’s multi- comparison test (*p<0.05, **p<0.01, ***p<0.001, ****p<0.0001; ns = non-significant). **E:** Spearman correlation analysis (two-tailed; 0.95% confidence interval) between PI4P and giantin fluorescence levels (CTCF) and also between each protein and dsRNA in RV-C15-infected HAE at 12hpi. **F and G:** 3D surface plot of RV-C15-infected HAE (at 12hpi) showing the immunodetection of dsRNA (red; **F and G**), giantin (green; **F**), PI4P (green; **G**); and colocalization (yellow) between dsRNA/giantin (**F**) or dsRNA/PI4P (**G**). **H and I**: Spatial colocalization analysis of dsRNA/giantin (**H**) and dsRNA/PI4P (**I**) in HAE infected with RV-C15, RV-A16, and RV-A2 at 12 hpi. The median colocalization ratio is noted on top of each bar. Statistical analysis was done by Kruskal-Wallis test following Dunn’s multi-comparison test.

In addition to a loss of Golgi integrity, we also observed enhanced detection of PI4P in cells with active RV replication compared to non-infected controls. Similar to giantin, PI4P was dispersed throughout the cytoplasm in infected cells (**Fig 3C**). To further assess the changes in giantin and PI4P levels, we compared the CTCF for these cellular targets in both non-infected and infected cells. Our data revealed both giantin and PI4P levels were elevated and strongly correlated in HAE infected with all RVs, though the change in PI4P was not significant for RV-A2 (**Fig 3D, 3E and S3 Fig**). A direct relationship was also observed between giantin and dsRNA CTCF in RV-C15- and RV-A16-infected cells while PI4P and viral dsRNA CTCF were only correlated in RV-C15 HAE (**Fig 3E, and S3 Fig**).

The close association between Golgi membranes and picornavirus replication has been shown by the colocalization between viral dsRNA and different Golgi markers (e.g., giantin, TGN- 46, GM130) [28, 31, 34]. Therefore, we next sought to evaluate the Golgi as a site for RV-C15 replication through colocalization analysis of dsRNA with giantin, and PI4P. Despite the strong correlation between giantin and PI4P observed during RV-infection (**Fig 3E and S3 Fig**), the ratio of colocalization between them was very low at pixel-intensity and spatial levels (**S1-3 Tables**). Surprisingly, the ratios of viral dsRNA/giantin and dsRNA/PI4P colocalization were also low using pixel-intensity based methods (**S4-9 Tables**). The 3D surface plot from RV-C15-infected cells at 12 hpi also demonstrated little evidence of colocalization between dsRNA and giantin (**Fig 3F**) or PI4P (**Fig 3G**). Spatial colocalization analysis confirmed these observations, with less than 9% of the total dsRNA detected at the same location as giantin or PI4P in RV-C15-infected cells (**Fig 3H and 3I, S4-9 Tables**). Thus, despite the fragmentation of the Golgi stacks and increase of PI4P in airway epithelial cells with active RV-C15 replication, our data indicate the Golgi is not the main site for viral genome replication and PI4P is not a marker for sites of RV-C15 replication in HAE as observed for other RVs [31, 34].

**Figure 4.**
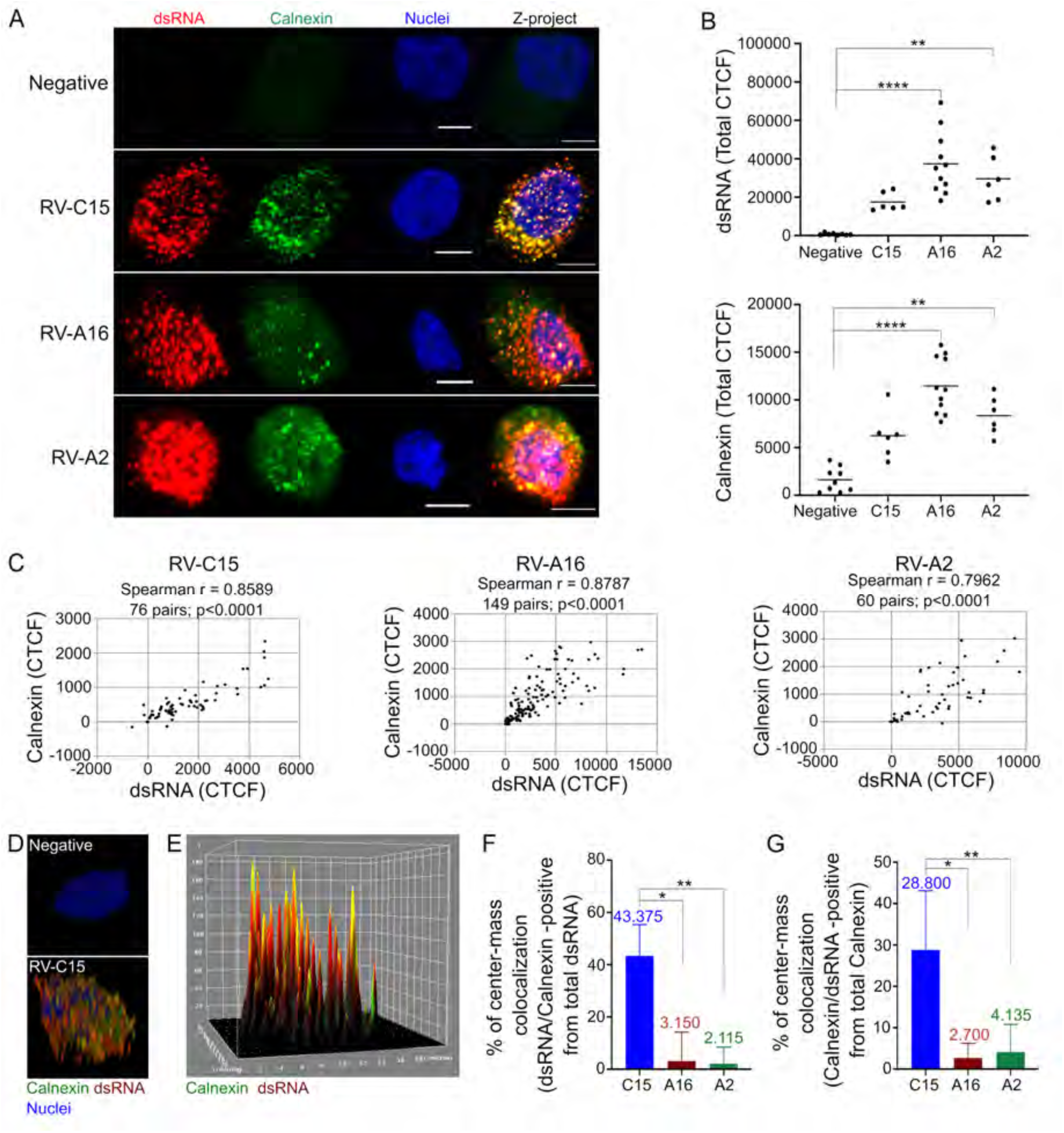
ER is a site for RV-C15 genome replication in HAE. **A**: Detection of calnexin (ER marker, green; nuclei, blue) in non-infected or rhinovirus-infected (dsRNA+, red) HAE at 12 hpi (z-stacks at 1µm of thickness; scale bar = 5µm). **B:** Quantification of fluorescence levels (CTCF; line = mean) of dsRNA and calnexin in non-infected cells and RV-infected (dsRNA+) HAE detected by immunofluorescence at 12 hpi. Statistical analysis was done using Kruskal-Wallis test following Dunn’s multi-comparison test (**p<0.01, **** p<0.0001). **C:** Spearman correlation analysis (two-tailed; 0.95% confidence interval) between calnexin and dsRNA fluorescence levels (CTCF) in RV-C15, -A16, or -A2-infected HAE at 12 hpi. **D:** 3D view of RV-C15-infected HAE shows the immunodetection of calnexin (green), viral dsRNA (red), and calnexin/dsRNA colocalization (yellow and orange) at 12 hpi (z-stacks at 1µm of thickness). **E:** 3D surface plot of RV-C15-infected HAE shows the detection of calnexin (green), dsRNA (red), and calnexin/dsRNA colocalization (yellow) by immunofluorescence at 12 hpi. **F-G:** Spatial colocalization analysis done in RV-C15, RV-A16, or RV-A2-infected HAE (dsRNA+) shows the ratio of dsRNA/calnexin (**F**) and calnexin/dsRNA (**G**) colocalization at 12 hpi. The median colocalization ratio is plotted on top of each bar; statistical analysis was done by Kruskal-Wallis test following Dunn’s multi-comparison test (*p<0.05; ** p<0.01).

### 4. Endoplasmic reticulum is a site for RV-C15 genome replication, but not RV-A16 or RV-A2

Given the low ratio of colocalization between RV-C15 dsRNA and Golgi markers, we next evaluated ER membranes as a potential site for viral genome replication. WB analysis of the ER-associated protein calnexin in RV-C15-infected HAE did not indicate a significant change compared to non-infected cultures at 12, 24, or 48 hpi (**S4 Fig**). However, at the single-cell level, calnexin fluorescence was not only elevated in cells infected with RV-C15, RV-A16, and RV-A2 compared to the negative control at 12 hpi (**Fig 4A and 4B**) but also directly correlated with fluorescence levels of viral dsRNA (**Fig 4C**). Interestingly, the 3D view of HAE infected with RV- C15 at 12 hpi showed calnexin spread throughout the cytoplasm, and there was a strong indication of calnexin and viral dsRNA colocalization (**Fig 4D**), which was further supported by 3D surface plot data (**Fig 4E**).

Pixel-intensity-based analysis confirmed the high level of colocalization between viral dsRNA and calnexin in cells infected not only with RV-C15 (56.3%), but also RV-A16 (28%) and RV-A2 (49.1%) at 12 hpi (**S10-12 Tables**). However, this high colocalization ratio was only confirmed for RV-C15-infected cells at the 3D level (spatial-colocalization analysis; **Fig 6F and 6G**), suggesting that, unlike RV-A16 and RV-A2, the ER is the main site for RV-C15 genome replication.

### 5. RV-C15 infection induces incomplete autophagy

Previous studies have shown picornaviruses, including several rhinoviruses, manipulate the autophagy pathway to mediate non-lytic release of progeny following replication [37]. Since the induction of autophagy is rhinovirus genotype-specific [37–39], we sought to determine if RV- C15 triggered the induction of autophagy in HAE. Towards this goal, we inoculated cultures with RV-C15 alongside RV-A16 and RV-A2 as controls, and probed for Lysosome-associated membrane glycoprotein 1 (Lamp-1; a lysosome marker) and LC3b (an autophagosome marker) by WB. LC3b protein levels were minimal in non-infected HAE, suggesting a low basal rate of autophagy while Lamp-1, LC3b-I, and LC3b-II expression varied across donors and time points during RV-C15 infection. Still, as these proteins were typically elevated at 12 hpi (**Fig 6A**), we further evaluated levels of Lamp-1 and LC3b in HAE by IF at this time point. Lamp-1 and LC3b detection in HAE was significantly stronger in RV-C15- and RV-A2-infected cells while the level of detection in RV-A16-infected HAE was similar to the negative control (**Fig 6B and 6C**). Curiously, the 3D view of non-infected cells revealed an apical localization of Lamp-1 while this protein was spread through the cytoplasm in RV-infected cells (**Fig 6B**). A high correlation ratio between Lamp1 fluorescence levels and dsRNA was obtained in RV-infected cells; however, LC3b levels strongly correlated with dsRNA only in cultures infected with RV-C15 and RV-A2 (**Fig 6D and S5 Fig**). Together, these data suggest RV-C15 and RV-A2 (but not RV-A16) induce autophagy to significant levels over the baseline in HAE.

**Figure 5.**
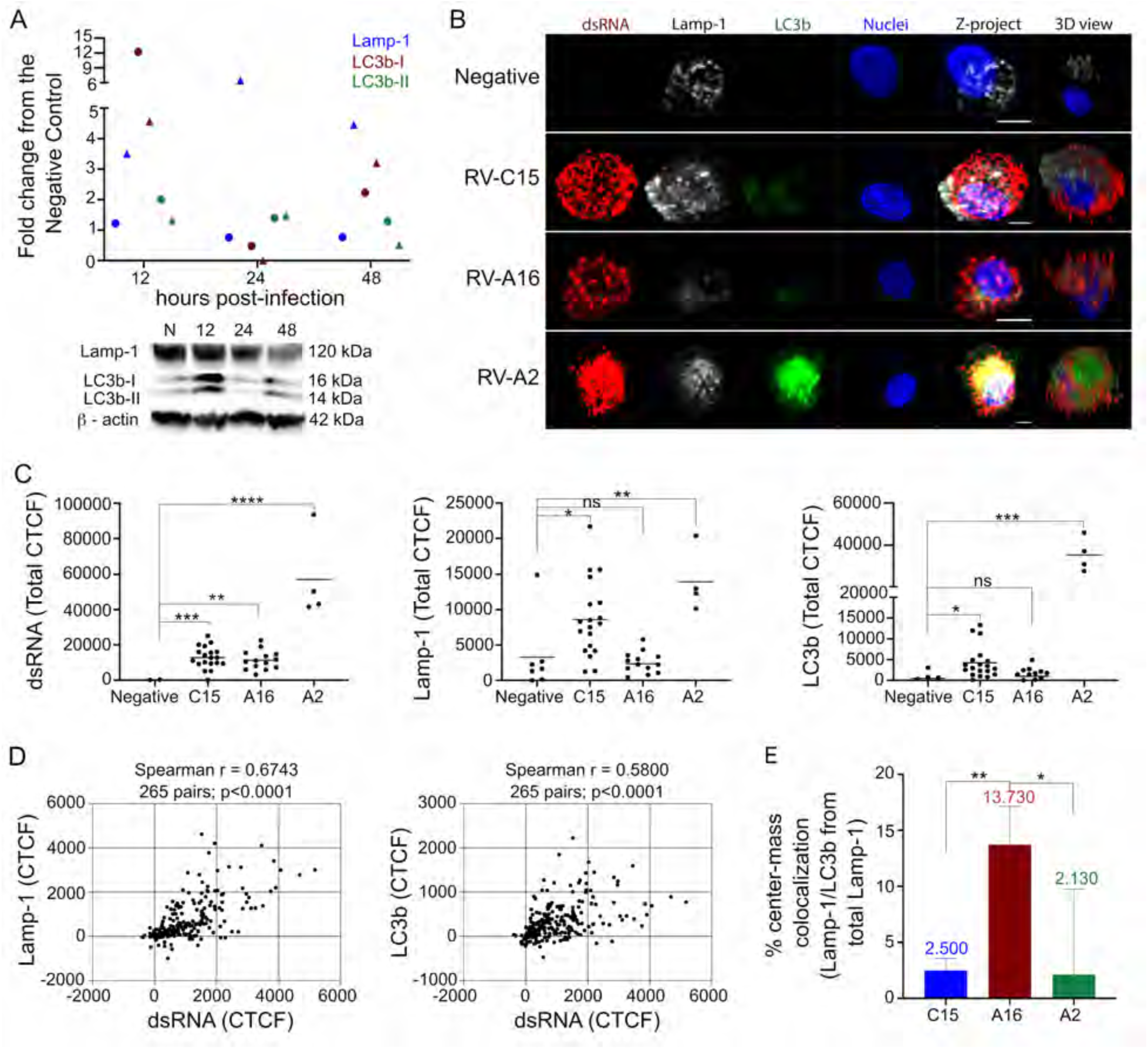
RV-C15 replication induces incomplete autophagy in HAE at 12 hpi. **A:** Fold change graph represents Lamp1, LC3b-I, and LC3b-II protein levels normalized to the endogenous control (actin) and compared to non-infected cultures. Data shown are from two independent donors, represented by circles and triangles. Blot below is from the donor represented by circles. **B:** Detection of Lamp-1 (gray), LC3b (green) and dsRNA (red; nuclei, blue) in non-infected HAE and HAE infected with RV-C15, RV-A16, or RV-A2 by immunofluorescence at 12 hpi (z-stacks at 1µm of thickness; scale bar = 5µm). **C:** Quantification of fluorescence levels (CTCF) for Lamp-1, LC3b, and dsRNA in non-infected and RV-infected (dsRNA+) HAE at 12 hpi. Statistical analysis was done by Kruskal-Wallis test following Dunn’s multi-comparison test (*p<0.05, **p<0.01, ***p<0.001, ****p<0.0001, ns = non-significant). **D:** Spearman correlation analysis (two-tailed; 0.95% confidence interval) between fluorescence levels for dsRNA and Lamp-1 or LC3b in RV-C15-infected HAE at 12 hpi. **E:** Spatial colocalization analysis of Lamp- 1/LC3b colocalization in RV-C15, RV-A16 and RV-A2 -infected (dsRNA+) HAE at 12hpi. The median colocalization ratio value is plotted on top of each bar; statistical analysis done by Kruskal- Wallis test following Dunn’s multi-comparison test (*p<0.05; ** p<0.01).

**Figure 6.**
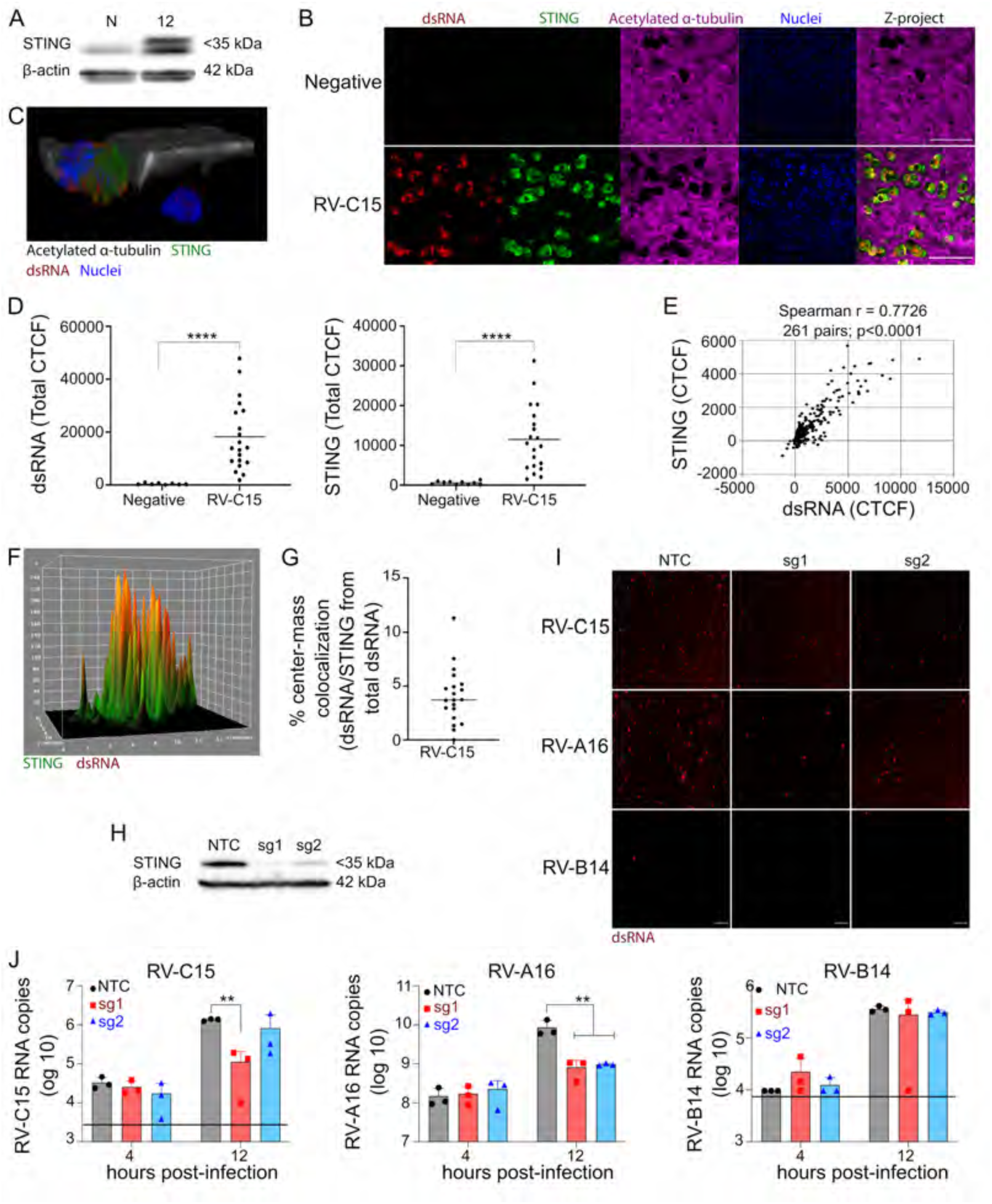
STING expression promotes RV-C15 replication in HAE at 12 hpi. **A:** Western blot showing STING expression at 48hpi in non-infected or RV-C15-infected HAE. **B:** Immunofluorescence detection of STING (green) in non-infected or RV-C15-infected (dsRNA; red) ciliated cells (acetylated α-tubulin, magenta) at 12 hpi (scale bar = 50µm). **C:** 3D view of RV-C15- infected HAE shows the detection of STING (green) and viral dsRNA (red) in ciliated cells (acetylated α-tubulin, gray) at 12 hpi (z-stacks at 1µm of thickness). **D:** Quantification of dsRNA and STING fluorescence levels (CTCF) in non-infected and RV-C15-infected (dsRNA+) HAE at 12 hpi. Statistical analysis was done by Mann-Whitney test (two-tailed; 0.95% confidence interval; ****p<0.0001). **E:** Spearman correlation analysis (two-tailed; 0.95% confidence interval) between fluorescence levels for dsRNA and STING in RV-C15-infected HAE at 12 hpi. **F:** 3D surface plot of RV-C15-infected HAE shows the detection of STING (green), dsRNA (red), and STING/dsRNA colocalization (yellow and orange) by immunofluorescence at 12 hpi. **G:** Spatial colocalization analysis shows the ratio of dsRNA/STING colocalization in RV-C15-infected HAE at 12 hpi (bar. = median). **H:** Western blot showing the STING expression in HAE derived from BCi-NS1.1 cells transduced with non-targeting control (NTC), sgRNA1 (sg1), or sgRNA2 (sg2). **I:** Immunofluorescence detection of dsRNA (red) in HAE derived from BCi-NS1.1 cells transduced with NTC, sgRNA1 (sg1), or sgRNA2 (sg2) infected with RV-C15, RV-A16, or RV-B14 at 12 hpi (scale bar = 100µm). **J:** qPCR to detect RV RNA in HAE derived from BCi-NS1.1 cells transduced with NTC, sgRNA1 (sg1), or sgRNA2 (sg2) and infected with RV-C15, RV-A16, or RV-B14 at 4 and 12 hpi. The black line represents the limit of detection (RV-C15 = 10^3.5^; RV-A16 = 10^5.9^ (not visible on the graph); RV-B14=10^4.8^); statistical analysis was done by Ordinary One-way ANOVA and Tukey’s multiple comparisons test (**p<0.01).

Since the inhibition of autolysosome formation at the termination of the autophagy pathway is characteristic of picornaviruses that induce autophagy to promote virus release [40], we further evaluated autolysosome formation defined by Lamp-1 and LC3b colocalization. Results from spatial colocalization analysis in cultures infected with RVs indicated greater detection autolysosomes in RV-A16-infected cells compared to RV-C15- and RV-A2-infected cells (**Fig 6E and S13-15 Tables**). Thus, these data suggest that, similar to RV-A2 but unlike RV-A16, RV-C15 induces incomplete autophagy in HAE.

### 6. STING expression promotes replication of RV-C15 in HAE

In this study, we observed RV-C15 genome replication in HAE to be associated with the ER (**Fig 4**) and the induction of incomplete autophagy at 12 hpi (**Fig 5**). Interestingly, STING, which localizes to the ER, was recently shown to induce the autophagy pathway during RV-A16 infection [41] and was identified as a proviral factor for RV-A and RV-C, but not RV-B, where STING overexpression facilitated viral genome replication in Huh-7 cells [24]. Based on these data, we investigated the expression of STING in HAE infected with RV-C15. Interestingly, we found elevated protein levels and the appearance of a second higher molecular weight band at 48 hpi compared to non-infected HAE (**Fig 6A**) as well as robust detection by IF in dsRNA(+) ciliated cells at 12 hpi (**Fig 6B**). Notably, 3D view analysis revealed that dsRNA/STING-double positive cells also exhibited dispersed distribution of acetylated alpha-tubulin, suggesting STING levels increase as replication progresses (**Fig 6C**). Indeed, the CTCF of STING not only increased significantly in HAE infected with RV-C15 (**Fig 6D**), but also strongly correlated with dsRNA levels at 12 hpi (**Fig 6E**). However, despite the 3D surface plot and pixel-intensity colocalization analysis suggesting high level of dsRNA/STING colocalization in RV-C15-infected HAE (**Fig 6F and S16 Table**), spatial colocalization analysis indicated the median colocalization ratio between both targets was lower than 5% (**Fig 6G**).

To better understand the importance of STING to RV-C15 genome replication, we genetically knocked-out STING using a CRISPR/Cas9 approach in immortalized human airway epithelial cells (BCi-NS1.1 cells [42]), differentiated them into HAE cultures (**Fig 6H**), and infected them with RV-C15, RV-A16, or RV-B14. Immunofluorescence detection of dsRNA at 12 hpi confirmed the susceptibility of control cultures expressing a non-targeting guide RNA to RV-C15, RV-A16, and RV-B14 infection (**Fig 6I**). However, the frequency of dsRNA (+) cells was lower in STING-depleted HAE infected with RV-C15 and RV-A16 (**Fig 6I**). While infection with RV-B14 was less efficient overall, a similar frequency of dsRNA+ cells were observed in both control and knockout HAE (**Fig 6I**). STING-knockout cultures also had significantly lower intracellular viral RNA copy numbers for RV-C15 and RV-A16 but not RV-B14, further validating our results (**Fig 6J**) and suggesting that the increase of STING expression in RV-C15-infected HAE has an advantageous impact on viral genome replication.

### 7. RV-C15 infection alters epithelial permeability, disposition of tight junction-associated proteins, and mucociliary clearance

Our results to this point highlight RV-C15-induced cytopathic effects at the single-cell level in HAE (**Fig 1-5**). To extend these observations, we quantified extracellular lactate dehydrogenase (LDH) at the culture-level as an indication of plasma membrane leakage. LDH release was restricted to the apical compartment and did not exceed 20% of the maximum levels obtained from lysed control cultures; however, LDH levels increased over time (**Fig 7A**) in line with RV-C15 replication kinetics (**Fig 1A and 1B**). Thus, we sought to further evaluate the global impact of RV-C15 infection on epithelial integrity. Interrogation of HAE permeability during RV- C15 infection revealed a significant, albeit transient, decrease in transepithelial electrical resistance (TEER) at 12 hpi (**Fig 7B**). Since the transepithelial transport of ions is mediated by pores formed by integral membrane proteins termed claudins [43], we next investigated the expression and localization of claudin-1 during RV-C15 infection in HAE. While claudin-1 protein levels in cultures infected with RV-C15 did not vary (**Fig 7C)**, we observed a change in claudin-1 distribution in RV-C15-infected cells at 12 hpi by IF (**Fig 7D**). To determine if this translocation was specific to claudin-1, we characterized Zona occludens 1 (ZO-1), a cytoplasmic tight junction-associated protein. Similar to claudin-1, ZO-1 protein levels for both subunits (ZO-1 +α and ZO-1 -α) did not vary in RV-C15-infected HAE compared to the negative control (**Fig 7C**), while ZO-1 disposition was more widespread in dsRNA(+) cells (**Fig 7E**). To quantify the effect of RV-C15 infection on ZO-1 disposition over time, we utilized the Junction Analyzer Program (JAnaP) [44–46], which allowed us to assess the profile of ZO-1 detected in non-infected and RV-C15-infected cells (**Fig 7F**). The increase of discontinuous ZO-1 quantified in this assay (**Fig 7G**) indicates the ZO-1 translocation in HAE (**Fig 7E**) parallels the progression of RV-C15-infection. Additionally, two profiles of discontinuous ZO-1 were evaluated, and ZO-1 was found in a more perpendicular than punctual disposition in RV-C15- infected HAE (**Fig 7H and 7I**).

**Figure 7.**
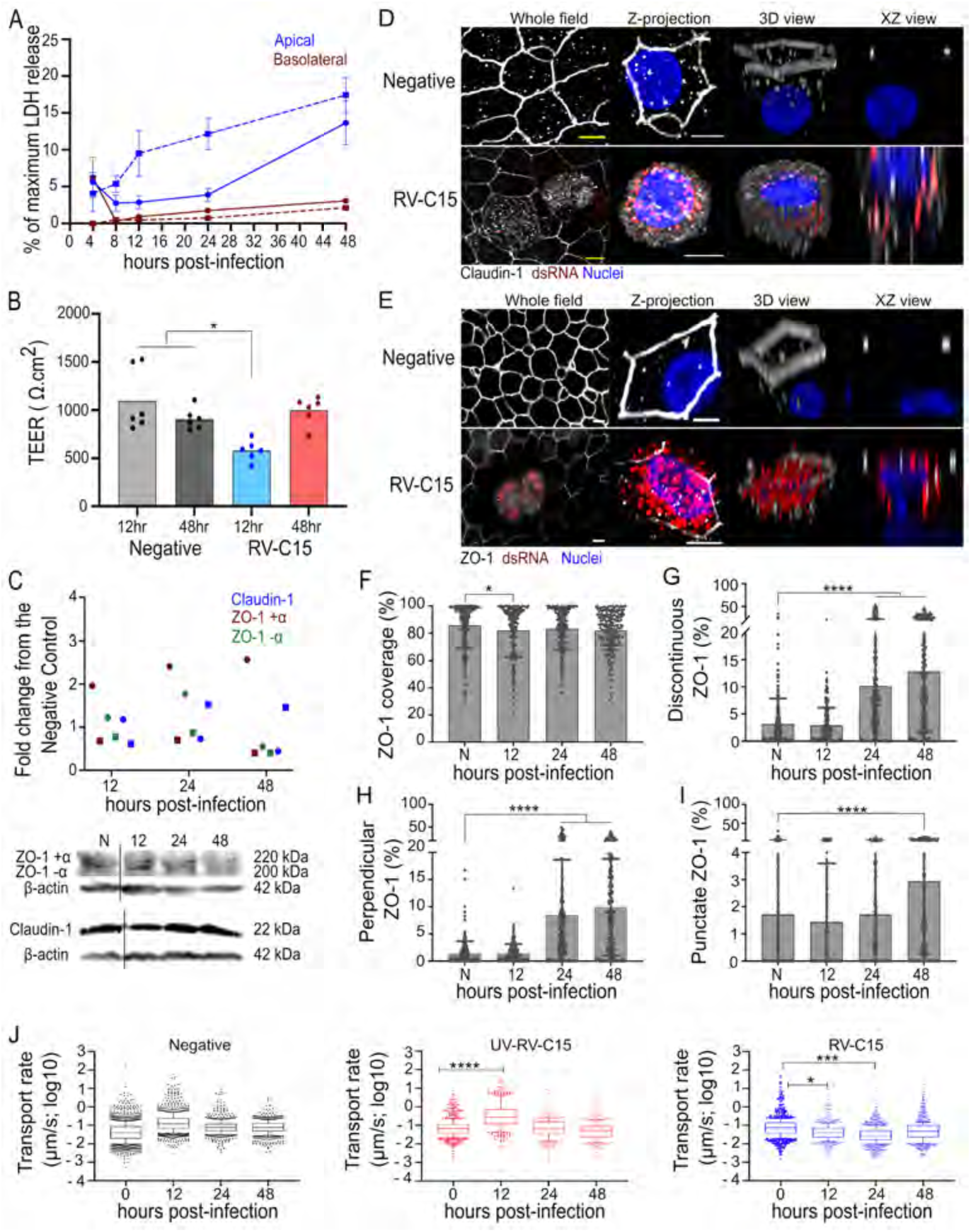
RV-C15 infection promotes translocation of tight junction proteins and impairs mucociliary clearance (MCC). **A:** Percentage of maximum LDH release graph shows the cytotoxicity of RV-C15 infection in HAE from two donors (continuous and dash lines). **B:** Quantification of transepithelial electrical resistance (TEER) in non-infected and RV-C15-infected HAE (two donors; each performed in triplicate). Statistical analysis was done by Ordinary one- way ANOVA following Tukey’s multiple comparisons test (*p<0.05). **C**: Fold change graph represents claudin-1 and zona occludens-1 (ZO-1) α+ and α- protein levels normalized to the endogenous control (actin) and compared to non-infected cultures. Data shown are from two independent donors, represented by circles and squares. Blot below is from the donor represented by squares; black line represents lanes not included in this analysis. **D-E:** Visualization of claudin-1 (**D,** gray) and ZO-1 (**E,** gray) in non-infected and RV-C15-infected (dsRNA+, red) HAE (nuclei, blue) detected by immunofluorescence at 12 hpi (z-stacks at 1µm of thickness; scale bar = 5µm). **F-I:** Quantification of ZO-1, detected by immunofluorescence, in non- infected and RV-C15-infected using the JAnaP software [44–46]. Total ZO-1 detected per cell perimeter (percentage of coverage; **F**); detection of discontinuous ZO-1 (disrupted; **G**) -perpendicular (**H**) and punctate (**I)** profiles. Statistical analysis was done by Ordinary one-way ANOVA following Dunnett’s multiple-comparison test (*p<0.05, ****p<0.0001). **J:** Mucociliary clearance (MCC) quantified in non-infected HAE (negative control) and HAE inoculated with either UV-RV-C15 or RV-C15 (10^10^ RNA). Statistical analysis was done by Ordinary one-way ANOVA following Dunnett’s multiple-comparison test (*p<0.05, ***p<0.001, ****p<0.0001).

Given the ciliated cell tropism RV-C15 and the important role of these cells in promoting mucociliary clearance (MCC), we also assessed the global impact of RV-C15 infection on mucus transport. HAE cultures were equilibrated at 34°C and the transport rate of red-fluorescent microspheres in the extracellular mucus gel was calculated immediately prior to inoculation, and up to 48 hpi. IF confirmed viral replication at 12 hpi in HAE inoculated with viable, but not UV-inactivated, RV-C15 (**S6 Fig**). The transport of microspheres indicated a slight increase in MCC in HAE inoculated with PBS and a similar, albeit significant, elevation in microsphere transport rate in UV-inactivated RV-C15-inoculated cultures at 12 hpi (**Fig 7J**), likely attributable to the addition of fluid on the apical surface after the T=0 time point. Despite this, RV-C15 infection resulted in a significant decrease in MCC at both 12 and 24 hpi (**Fig 7J**) indicating viral replication impairs this innate host defense mechanism.

## Discussion

Rhinovirus is becoming increasingly recognized as a cause of both upper and lower respiratory tract infection [4–8]. Furthermore, the risk for development of asthma in children after infection by RV, especially RV-C [8, 13, 14], highlights the clinical relevance of these viruses. Due to restricted receptor expression in traditional immortalized cell culture systems [26], we investigated RV-C15 replication using an *in vitro* model of HAE that supports the entire RV-C life cycle. Previous studies in similar systems have shown the impact of temperature and cell differentiation stage on RV-C replication [22, 25, 27]. Here, we detail a multi-step replication curve for RV-C15 (**Fig 1A and 1B)**, indicating the peak of intracellular viral load at 12 hpi by qPCR and IF (**Fig 1A and 1B, S1A Fig**) and peak viral release at 24 hpi (**Fig 1A and 1B**) in agreement with a previous report characterizing RV-C11 and RV-C15 infection in bronchial cells [26].

In well-differentiated HAE, we detected CDHR3 on the apical surface of ciliated cells (**Fig 1C-1E**). Interestingly, we did not observe any evidence of cytosolic CDHR3 in non-infected cells, differing from previous immunofluorescence data [21, 23] but in agreement with CDHR3 detection on cilia by TEM [21]. However, CDHR3 distribution changed, and fluorescence levels decreased, in RV-C15-infected cells (**Fig 1C-1E, and 1G**) without altering levels of acetylated alpha-tubulin or FoxJ1 (**S1G and S1H Fig**). Unlike ICAM-1 and LDLR (receptors for RV-A and RV-B, respectively), the dynamics of CDHR3 during infection and associated pathway(s) of RV-C entry remain unknown. After endocytosis of RV-A and RV-B particles, ICAM-1 and LDLR are either degraded in the lysosome with empty capsids, or LDLR is trafficked from the endosome to be recycled [47]. The intracellular distribution of CDHR3 after RV-C15 infection in HAE suggests CDHR3 follows the same path as other RV receptors. In contrast, global expression of CDHR3 protein quantified by WB increased 4-fold at 12 hpi in RV-C15 infected epithelium (**S1F Fig)**. Since the function of CDHR3 is unknown, the trigger for this elevated expression and what impact this may have on infection remains unclear. Nonetheless, similar up-regulation was detected for ICAM-1 and LDLR in primary cell cultures and human volunteers after RV infection [48–50].

After confirming the susceptibility of our HAE models to RV-C15 infection, we further investigated the localization and association of replication complexes with various organelle markers at the single-cell level. Using dsRNA as a marker for virus replication, we observed both perinuclear and “ring-like” distribution of replication centers closer to the basolateral membrane during RV-C15, RV-A16, and RV-A2 replication (**Fig 2A-2C and S2A-S2D Fig)**. To our knowledge, the latter profile has not been described during RV or any other picornavirus infection, and our TEM data indicates the formation of large intracellular vesicles filled with unknown electron-dense content may be the underlying cause of altered dsRNA disposition. Whether these large vesicles play a specific role in RV replication or if they are simply indicative of cellular changes that precede cell death is unknown.

Notably, our TEM analysis also identified smaller, double-membraned vesicles, similar to those identified during replication of other RVs and related picornaviruses [28–31], that likely represent sites of RV-C15 genome replication (**Fig 2D and 2E**). Prior reports identified the induction of replicative complexes derived from the ER by the poliovirus 2BC and 3A proteins [51, 52] and these same proteins in RV-A16 were associated with the ER at an early stage of viral infection [35]. Nonetheless, work to date suggests the Golgi is the primary site for picornavirus genome replication [28, 31, 34]. Further, the formation of replicative complexes has been associated with cholesterol exchange driven by OSBP1 [31], accumulation of ER-vesicles resulting from the inhibition of ERGIC-to-Golgi transport [36], and fragmentation of Golgi stacks induced by the viral 3A protein [34, 35]. In this study, we observed an increase of giantin and PI4P levels, dissolution of Golgi stacks, and spread of PI4P-positive vesicles in RV-C15-infected cells (**Fig 3A-D)** suggesting RV-C impacts Golgi membranes similar to other picornaviruses. However, despite the strong correlation between dsRNA and fluorescence levels of both giantin and PI4P (**Fig 3E**), colocalization analysis indicated neither giantin- nor PI4P-positive vesicles are the main site for RV-C15 genome replication (**Fig 3H and-3I**). Similar observations were made in RV-A16- infected HAE (**Fig 3**) although dsRNA fluorescence levels only strongly correlated with giantin and not PI4P (**S3A Fig)**. These data were unexpected, as prior work in HeLa and nasal epithelial cells noted PI4P enrichment of Golgi membranes during RV-A16 genome replication [31]. Different from the literature [31], Golgi and PI4P-positive vesicles were not the main site for RV- A2 genome replication in HAE either (**Fig 3H and 3I**), and neither the significant increase in levels of giantin nor insignificant increase in PI4P correlated strongly with viral dsRNA (**Fig 3D and S3B Fig**).

Most notably, our results identified the ER as the main site for RV-C15 replication through elevated levels of calnexin in RV-infected HAE (**Fig 4A and 4B**) and a high ratio of dsRNA/calnexin colocalization in cells infected with RV-C15, but not RV-A2 and RV-A16 (**Fig 4E- G**). The increase in calnexin levels observed in this study could be the result of ER stress in infected cells, as reported for RV-A1B and RV-A16 [53, 54]. The RV-A16 2B protein [54] forms pores on ER membranes, promoting calcium ion efflux, which increases membrane permeability and triggers stress. These effects on ER membranes have been associated with more efficient release of vesicles that may represent additional sites for picornavirus replication [55]. Notably, inhibition of ER stress resulted in a decrease in RV-A1B replication [53], corroborating the hypothesis that ER-derived vesicles are a site for viral replication [55] and supporting our conclusion that RV-C15 replication is associated with the ER in HAE (**Fig 4E-G**).

In the final stages of the viral life cycle, progeny picornavirus virions are released through cell lysis or by usurping the autophagy pathway [37–39]. Exploitation of the autophagy pathway was originally shown for poliovirus [40] and is characterized by virus-mediated inhibition of autolysosome formation and release of autophagosomes full of nascent virions that likely protect particles from the immune response [38]. As expected [39], RV-A2 induced autophagy in HAE in contrast to RV-A16 (**Fig 5B and 5C**) based on elevated Lamp1 and LC3b levels. However, a very low ratio of Lamp-1/Lc3B colocalization indicated a failure to form autolysosomes (**Fig 5E**). The results obtained in RV-C15-infected HAE were similar to RV-A2, indicating the induction and manipulation of the autophagy pathway (**Fig 5B, 5C, and 5E**). To our knowledge, this is the first time the autophagy pathway has been analyzed during RV-C infection.

The identification of ER as a site for genome replication and induction of autophagy during RV-C15 infection of HAE led us to investigate STING, a protein that has been associated with induction of ER stress and autophagy [41, 56]. In this study, we detected an increase in STING expression in RV-C15-infected HAE (**Fig 6A, 6B, and 6D**) which correlated with levels of dsRNA (**Fig 6E**). However, this correlation was not related to a close interaction between STING and dsRNA, as the median colocalization ratio at the spatial level was lower than 5% (**Fig 6G**). Additionally, the absence of STING impacted viral genome replication, resulting in significantly lower intracellular viral RNA levels at 12 hpi for RV-C15 and RV-A16, but not RV-B14 (**Fig 6I and 6J**). These data, which assayed endogenous STING expression in RV-infected HAE, corroborate published data in undifferentiated cells indicating STING expression is important for RV-A and RV-C genome viral replication [24]. STING is a ER-resident transmembrane protein which can be phosphorylated by TANK-binding Kinase I (TBK1) or translocated to the ER-Golgi intermediate compartment (ERGIC) before phosphorylation [57]. The phosphorylation of STING in the ER is necessary for autophagy induction during RV-A16 infection [41], while the translocation to the ERGIC is required for ER stress [56]. The low ratio of dsRNA/STING spatial colocalization at 12 hpi (**Fig 6G**) is surprising and indicates that either genome replication of RV-C15 occurs at different sites on ER membranes (STING-negative) or most of STING had already been translocated to ERGIC. RV-C15 replication also induced autophagy in HAE, though more experiments are needed to relate this phenomenon to STING activation.

Given the observed impacts of RV-C15 replication in infected cells, we further probed the effects of viral infection on epithelial integrity and functionality. The level of cytotoxicity of RV-C15 infection in HAE, measured by LDH release, increased overtime in the apical compartment only (**Fig 7A**), in agreement with viral cellular tropism for ciliated epithelial cells that face the luminal surface of the airway **(Fig 1**) [22, 25]. The dissociation of tight junctions during viral infection, including RV-A and RV-B, has been associated with loss of epithelial-selective permeability and increase in bacteria translocation across the epithelium [58]. A previous analysis of RV-A16, RV- A1B, and RV-A39 found the reduction of TEER and increase in epithelium permeability for inulin were associated with both claudin-1 and ZO-1 dissociation [58]. Similar to these reports, RV-C15 temporarily increased epithelial permeability to small particles and ions (as indicated by a temporary loss in TEER, **Fig 7B**) which may be due, at least in part, to claudin-1 translocation (**Fig 7C and 7D**). Additional claudins (e.g., claudin-3, -4 and -5) that are also expressed in the airway epithelium and known to impact transcellular transport were not assayed here [59, 60]. Still, the altered expression of tight junction proteins was not restricted to claudin-1 as we also detected ZO-1 throughout the cytoplasm in dsRNA(+) cells without any change in overall protein levels during infection (**Fig 7C and 7E**). Despite being a tight junction protein, ZO-1 interactions are more important to cell polarization and paracellular permeability [61]. Although we did not check paracellular transport of large particles, disrupted states for ZO-1 that indicated progressive translocation with ongoing viral infection (**Fig 7F-I**) could impact cellular organization. Nonetheless, the loss of TEER and changes in claudin-1 and ZO-1 observed in this study for RV- C15 were less dramatic than previous reports for RV-A and RV-B [37, 39, 58].

Beyond epithelial barrier integrity, we also found RV-C15 infection impacts mucociliary clearance functionality. Similar to a previous study [62], MCC decreased significantly after 12 hpi in HAE infected with RV-C15 but not UV-inactivated virus, indicating that active viral replication interferes with this cellular mechanism of pathogen removal (**Fig 7J**). We speculate that the temporary increase in MCC observed in cultures inoculated with UV-inactivated virus is due to the addition of fluid on the apical surface. However, this increase was not significant for the negative control and therefore may indicate that the host response to incoming viral particles stimulates MCC. The coordinated beating of ciliated cells drives the basal MCC rate of ∼5.5 mm/min, which can be altered by mucus composition, temperature, and humidity [63–65]. Indeed, mucus hypersecretion [66] and changes in mucus composition [67] are both associated with RV infections. In this study, we noted altered distribution of acetylated alpha-tubulin in cells with active viral replication (**Fig 1D and 1F**) which likely indicates impaired cilium structure and function in these cells. Whether the progressive loss of cilia function and eventual death of infected cells contributes to our observed change in MCC during RV-C15 infection is not clear.

In conclusion, our present study expands our current understanding of RV replication in a physiologically-relevant setting, substantiating previous observations related to virus-induced membrane reorganization, and demonstrating that RV-C15 displays many typical features of picornavirus infection. In addition, we identify RV-C15 replication in association with ER membranes unlike previously characterized RVs. While we speculate that these data identify a unique feature of RV-C replication, our study is limited by the use of only one RV-C genotype; thus, further analysis to confirm these observations using additional RV-C isolates is important. Notwithstanding, our findings underscore the fact that virus-host interactions critical for rhinovirus replication are not always conserved, possibly contributing to their different clinical profiles, and supporting further investigation into this group of pathogens.

## Materials and Methods

### 1. Primary and immortalized cell culture and viral stocks

Human airway tracheobronchial epithelial cells isolated from airway specimens from donors without underlying lung disease were provided by Lonza, Inc. Primary cells derived from single patient sources were expanded on plastic and subsequently plated (5 x 10^4^ cells / well) on rat-tail collagen type 1-coated permeable transwell membrane supports (6.5 mm, #3470; Corning, Inc., Corning, NY). HAE cultures were grown in Pneumacult-Ex basal medium (#05008, StemCell Technologies, Seattle, WA), or Pneumacult-ALI medium (#05001, StemCell Technologies, Seattle, WA) with provision of an air-liquid interface for approximately 6 weeks to form differentiated, polarized cultures that resemble *in vivo* pseudostratified mucociliary epithelium.

H1 HeLa (#CRL-1988; ATCC, Manassas, VA) and HEK-293T (#CRL-11268; ATCC) cells were cultivated in Dulbecco’s Modified Eagle Medium (DMEM; #11965118; Gibco – Thermo Scientific, Waltham, MA) supplemented with fetal bovine serum (FBS; GenClone FBS; Genesee Scientific, San Diego, CA) and 1% penicillin/streptomycin (#15140122; Gibco). All cell cultures were maintained at 37°C with 5% CO_2_.

RV-C15, RV-A2, RV-B14, and RV-A16 were rescued from infectious clones kindly donated by Drs. James Gern and Yuri A. Bochkov, (pRV-C15), Dr. Wai-Ming Lee (pRV-A2) and Dr. Ann Palmenberg (pRV-B14 and pRV-A16) from University of Wisconsin – Madison. Rescue of infectious RV was done according to published protocols [68, 69] in H1 HeLa cells, with one modification: viral RNA transfection was performed using jetPRIME reagent (#114-07; Polyplus transfection, Illkirch, France).

### 2. CRISPR-Cas9-mediated knockout of STING in HAE

Single guide RNAs (sgRNA) targeting STING or no known target (NTC) and flanked by restriction sites for cloning into the pLentiCRISPRv2 backbone [69] with eGFP replacing puromycin selection were as follows: sgRNA1- 5’-CACCGCATATTACATCGGATATCTG-3’ and 5’-AAACCAGATATCCGATGTAATATGC-3’; sgRNA2- 5’- CACCGACTCTTCTGCCGGACACTTG-3’ and 5’-AAACCAAGTGTCCGGCAGAAGAGTC-3’; NTC- 5’-CACCGGCACTACCAGAGCTAACTCA-3’ and 5’- AAACTGAGTTAGCTCTGGTAGTGCC-3’. Lentiviral stocks were generated by co-transfection of 1μg pLentiCRISPRv2 (Addgene plasmid #52961 donated by Dr. Feng Zhang) [70], 0.2 μg pCMV-VSV G (Addgene plasmid #8454 donated by Dr. Bob Weinberg) [71]), and 0.7 μg psPAX2 (Addgene plasmid #12260 donated by Dr. Didier Trono) into HEK-293T cells with jetPRIME reagent (Polyplus transfection). Lentivirus-laden supernatant was collected and replaced at 24 hour intervals up to 72 hours, pooled, and filtered to remove viable cells and debris.

For target cell transduction, lentivirus-containing supernatants were applied to BCi-NS1.1 cells (kindly provided by Drs. Matthew Walters and Ronald Crystal [42]), maintained as HAE at 40-60% confluence with a final concentration of 20 mM HEPES (Gibco) and 4 μg/mL polybrene (Thermo Scientific). Cells were then centrifuged (1,000 x g for one hour at 37°C) and incubated at 37°C with 5% CO_2_ for 6 hr. The inoculum was removed and replaced with fresh Pneumacult-Ex Plus media (StemCell Technologies). At 60-80% confluence, eGFP-positive cells were enriched by fluorescence-activated cell sorting on a BD FACSAria-II Cell Sorter (BD Bioscience, San Jose, CA). These sorted BCi-NS1.1 cells were then expanded, seeded at 3.3x10^4^ cells/well on 6.5mm rat-tail collagen type I-coated transwell membranes, and cultured at air-liquid interface as previously described for HAE.

### 3. Titration of rhinoviruses by plaque assay and quantitative real-time PCR (qPCR)

RV-A2, RV-B14, and RV-A16 stocks were titrated by plaque assay method using H1 HeLa cells plated in 24-well dishes and a published protocol with modifications [72]. Briefly, 90% confluent H1 HeLa monolayers were inoculated with ten-fold serial dilutions of the viral stock in McCoy’s medium with 2% FBS and 30mM MgCl_2_. After 1 hour incubation at room temperature (RT), the inoculum was exchanged for a semi-solid overlay composed of 0.6% of bacteriological agar (#A5306; Sigma-Aldrich, Saint Louis, MO) in DMEM-F12 (#12500062; Gibco), supplemented with 1% FBS (Genesee), 1% penicillin/streptomycin (Gibco), 1% L-glutamine (#25030081; Gibco), and 30mM MgCl_2_ (Sigma-Aldrich). The plate was incubated at 34°C for 72 hours and the cells were fixed with 38% formaldehyde (#47608; Sigma-Aldrich) in a 0.15M saline solution. The fixing solution was removed together with the overlay after a 24-hour incubation at RT and plaques visualized by staining the monolayer with 0.1% crystal violet (#C0775; Sigma-Aldrich) diluted in a 20% ethanol (200 proof) solution.

RV-C15 was titrated by qPCR, which was also used to quantify RV-A16 and RV-B14 in experiments done in CRISPR/Cas9-modified HAE. To quantify viable viral particles, 10µl of RV- C15 stock or 100µL of experimental sample was first treated with 1u RNase A (#EN053; Thermo) and 1u DNase I (#18047019; Invitrogen – Thermo Scientific) for 1 hr at 37°C. Viral RNA was then extracted using a QIAamp Viral RNA Mini Kit (#52904; Qiagen, Hilden, Germany); total RNA was extracted from cells using a RNeasy Mini Kit (#74104; Qiagen). Next, up to 1µg of RNA was used in a reverse transcription reaction (High Capacity cDNA Reverse Transcriptase Kit; #4368814; Applied Biosystem – Thermo), with random hexamer primers, as per the manufacturer’s protocol. qPCR was done using 3µl of cDNA, 10pM of HRVTF63 forward (5’– ACMGTGTYCTAGCCTGCGTGGC–3’) and HRVTR reverse (5’–GAAACACGGACACCCAAAGT GT–3’) primers, 10pM HRVTF probe (5’–FAM/TCCTCCGGCCCCTGAAT/BHQ1–3’), and LuminoCT Taqman Master Mix (#L6669; Sigma-Aldrich) according to the manufacturer’s protocol. For absolute quantification, a standard curve was delineated using cDNA generated from ten-fold serial dilutions of *in vitro*-transcribed pRV-C15, pRV-A16, and pRV-B14.

### 4. RV-infection in HAE and H1 HeLa cells

One week before the experiment in HAE, cells were washed 3 x 30 minutes with 100µl of phosphate buffered saline (PBS; #P5119; Gibco) at 37°C to remove excessive mucus. In CRISPR/Cas9-modified BCi-NS1.1-derived HAE an additional wash was performed immediately before inoculation. HAE cultures were inoculated on the apical surface with 10µl of PBS (negative control), sucrose-purified RV-C15 (10^10^ copies of RNA), RV-A16 (5x10^5^ PFU), RV-B14 (5x10^5^ PFU), or RV-A2 (5x10^5^ PFU) and incubated at 34°C with 5% CO_2_ until sample collection. In the replication curve and cytotoxicity assays, the inoculum was removed, and the apical surface rinsed with PBS, at 4 hpi.

For HAE culture sample collection, 100µl of PBS (Gibco) was added to the apical chamber and harvested after a 30-minute incubation at 34°C; basolateral samples (500µl ALI media) were collected directly from underneath the culture. Cells were harvested from the transwell membranes using 350µl RLT buffer (#74104; Qiagen) for downstream RNA extraction, or 75µl of Radio Immuno-Precipitation Assay (RIPA) buffer (#89900S; Thermo) plus protease inhibitors (#88666; Thermo Scientific) for subsequent Western blot (WB) assays. The aliquots for RNA extraction and WB were stored at -80°C and -20°C, respectively. Alternatively, cultures were fixed with 4% (w/v) freshly-prepared paraformaldehyde (#157-8-100; Electron Microscopy Sciences (EMS), Hatfield, PA) for immunofluorescence (IF) assays or 2% (w/v) glutaraldehyde in 0.1M cacodylate buffer for downstream transmission electron microscopy analysis.

### 5. IF assays in HAE and H1 HeLa cells

HAE cultures were fixed with 100µl (apical) and 500µl (basolateral) 4% (w/v) freshly prepared paraformaldehyde (#157-8-100; EMS). After a 15-minute incubation at RT, the culture was washed once for 3 minutes with PBS and then incubated for 10 minutes at RT with a quenching solution (25mM NH_4_Cl diluted in PBS) before being washed twice for 3 minutes each with PBS. Permeabilization of cell membranes was done using 0.2% Triton-X diluted in PBS; after a 15-minute incubation at RT, the cells were washed 3 x 3 minutes with PBS. A blocking solution (10% normal donkey (#017-000-121) or normal goat serum (#005-000-121, Jackson ImmunoResearch Labs, West Grove, PA) diluted in PBS with 0.2% Tween-20) was added and only removed from the apical side after a 10-minute incubation at RT. The apical surface of the culture was washed 3 x 3 minutes with PBS. Primary antibody diluted in PBS with 0.2% Tween- 20 and 1% Bovine Serum Albumin (BSA; #BP9700100; Fisher Scientific, Hampton, NH) was added to the apical chamber and incubated overnight at 4°C, protected from light. The apical surface was washed 3 x 5 minutes with PBS and incubated with the secondary antibody diluted in PBS with 1% BSA. After a 2-hour incubation at RT, protected from light, the secondary antibody solution was removed and the apical surface of HAE was washed 3 x 5 minutes with PBS. If necessary, additional stains were done by repeating this protocol; reducing the secondary antibody incubation to 20 minutes when the next primary antibody applied was generated in the same species as another primary antibody used in the same experiment. Fluorescent-conjugated primary antibodies were applied during the final round of staining, where applicable. Nuclei were visualized in the final step by adding 1ng/ml of Hoechst 33342 (#H3570; Invitrogen, Thermo) diluted in PBS with 0.2% Tween-20 to the apical surface. After a 5-minute incubation at RT, the cultures were washed 3 x 5 minutes with PBS, after which the transwell membrane was separated from the plastic holder and mounted on a glass slide using VectaShield Antifade mounting medium (#H-1000-10; Vector Laboratories, Burlingame, CA) and 1.5mm-thick cover glass.

Images were acquired with a Zeiss Axio Observer 3 Inverted fluorescence microscope (Cell Observer HS image system, Zeiss Axiocam 503 mono camera, optimal acquisition mode, AIM-Zen 2007 software) equipped with EC-Plan-NEOFLUOR 20x/0.5NA Ph2 and 40x/0.75NA Ph2 air objectives. Higher resolution images and z-series optical sections (at 1µM intervals) were acquired with a LSM710 Zeiss confocal microscope (Argon laser, pinhole of 1 airy unit (AU), zoom factor of two, optimal acquisition mode, AIM-Zen 2009 software) equipped with a 63x/1.4NA Oil DIC Plan Apo objective at the Imaging Core, University of Maryland, College Park. All images were analyzed using Fiji – ImageJ v.2.1.0/1.53c software [73]. Z-series from a pre-selected region of interest (ROI) are displayed as maximum intensity z-projection, orthogonal views (XY, XZ and YZ planes), and 3D surface plots; they were also used for 3D volume reconstruction (3D view) using Fiji [73].

The IF protocol described above was used in all assays with the following exception: for PI4P detection, permeabilization was done using a 20mM Digitonin solution (diluted in PBS). Antibodies used in this study are listed in the tables below.

**Table 1:** Primary antibodies used in Immunofluorescence assays.

| Primary Antibody | Dilution | Catalog # | Source |
| --- | --- | --- | --- |
| Mouse IgG2a anti-dsRNA (J2) mAb | 1:1000 | #10010200 | SCICONS, Budapest, Hungary |
| Mouse IgG2b anti-acetylated alpha-tubulin (6-11B-1) mAb A647- conjugated | 1:50 | #sc-23950-AF647 | Santa Cruz Biotechnology<br>Incorporation, Dallas, Texas |
| Mouse IgG1 anti-FoxJ1 mAb (2A5) | 1:200 | #14-9965-82 | Invitrogen – Thermo |
| Rabbit IgG anti-claudin-1 mAb [EPRR18871] | 1:500 | #ab211737 | Abcam, Cambridge, UK |
| Goat IgG anti-calnexin polyclonal antibody | 1:500 | #PA5-19169 | Thermo |
| Rabbit IgG anti-giantin pAb - Golgi Marker | 1:500 | #ab80864 | Abcam |
| Rabbit IgG anti-cadherin-28 (CDHR3) pAb | 1:500 | #orb182906 | Biorbyt, Cambridge, UK |
| Rabbit IgG anti-Lamp1 (D3D11) mAb | 1:500 | #9091 | Cell Signaling, Danvers, MA |
| Rabbit IgG anti-LC3b (EPR18709) mAb A647- conjugated | 1:100 | #ab225383 | Abcam |
| Mouse IgM anti-PI(4)P (clone PI4-2) mAb | 1:200 | #Z-P004 | Echelon Bioscience, Salt Lake City, Utah |
| Rabbit IgG anti-TM173/STING polyclonal antibody | 1:100 | #19851-1AP | Proteintech, Rosemont, IL |

**Table 2:**
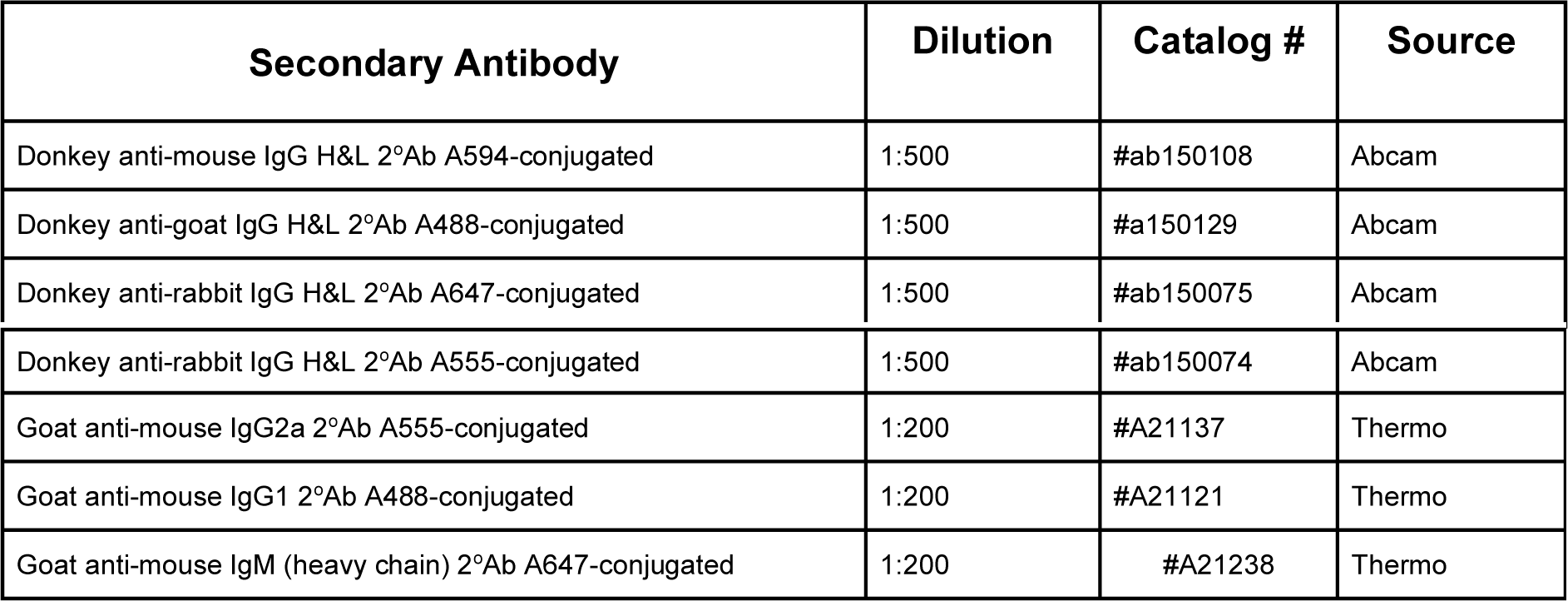
Secondary antibodies used in Immunofluorescence assays.

### 6. Quantification of fluorescence levels and colocalization analysis

The .CZI files from z-stacks collected with a step-size at 1µM were analyzed at region and single-cell levels using Fiji - ImageJ v.2.1.0/1.53c software [73]. For single-cell analysis, the selection of the region of interest (ROI) was based on positivity for dsRNA(+) in RV-infected HAE. At least 5 cells per region were selected to increase rigor.

Quantification of fluorescence levels was done in ROI single-cell images following the Corrected Total Cell Fluorescence (CTCF) method [74]. Total CTCF (equal to the sum of CTCF per slice / sample) was calculated per channel and statistically significant differences determined by Mann-Whitney U (Two-tailed; p<0.05 significance; 0.95% confidence interval) using Prism GraphPad v.9 software (GraphPad Software, San Diego, CA). Individual CTCF values were used in correlation analysis by the Spearman method (Two-tailed; p<0.05 significance; 0.95% confidence interval) using Prism GraphPad v.9 (GraphPad Software).

To evaluate the ratio of colocalization between two markers, the background was subtracted from ROI single-cell images following application of a threshold model and watershed filter to better identify the centroids. The final images were first used in colocalization analysis by pixel-intensity based methods (Pearson, threshold Manders, and Van Steensel’s methods), followed by spatial analysis based on the distance between centers of mass [75]. All colocalization analysis was done using the Just Another Co-localization Plug-in (JACoP) plugin from Fiji [75].

### 7. Transmission Electron Microscopy (TEM) in HAE

The TEM protocol used in this study was based on published methods [29] with modifications, as follows: Cultures were fixed with 2% (w/v) glutaraldehyde in 0.1M cacodylate buffer (100μl on top and 500μl on the bottom) for 60 minutes at RT. The transwell membranes were then separated from the plastic holder and transferred to a new 24-well plate where a second fixation step was carried out with 1% of osmium tetroxide (OsO_4_) in 0.1M cacodylate buffer plus 1% potassium ferricyanide (K_3_Fe(CN)_6_). Two percent uranyl acetate (diluted in distilled H_2_O) was used as post-fixative. The membrane was then incubated in propylene oxide before being embedded in Spurr’s Resin. The samples were sectioned to 60–90 nm with a diamond knife (DiATOME) and ultramicrotome (Reichart-Jung) and two slices were placed per copper grid (EMS). The images were obtained using a Hitachi S-4700 Field Emission Scanning Electron Microscope with transmitted electron detector in the Laboratory for Biological Ultrastructure, University of Maryland, College Park.

### 8. Immunoblotting assays

Total protein in cell lysates (stored at -20°C in RIPA buffer) were quantified by BCA Protein Assay (#23225; Pierce). A total of 20µg per sample (or 25µg and 30µg for STING and ZO-1 blots, respectively) was then separated on Novex WedgeWell 4-20% Tris-Glycine gels (#XP04202BOX; Invitrogen-Thermo) and transferred to PVDF membranes. Membranes were blocked with 5% non-fat milk solution (diluted in 0.1% Tween-20 in Tris-buffered saline (TBS-T)) or 3% BSA (diluted in TBS-T) for the detection of the tight junction proteins and incubated overnight at 4°C with the primary antibody. Following a series of washes in TBS-T, membranes were incubated for 1 hr at RT with the appropriate peroxidase-conjugated secondary antibody. All antibodies were diluted in 5% non-fat milk TBS-T solution. Blots were visualized using SuperSignal West Femto Maximum Sensitivity Substrate (#34094; Thermo) or SuperSignal West Dura Extended Duration Substrate (#34075; Thermo).

**Table 3:** Primary and secondary antibodies used in immunoblotting assays.

| Primary Antibody | Dilution | Catalog # | Source |
| --- | --- | --- | --- |
| Mouse IgG2b anti-acetylated alpha-tubulin (acetyl K40) monoclonal antibody | 1:5000 | #ab24610 | Abcam |
| Mouse IgG1 anti-FoxJ1 monoclonal antibody (2A5) | 1:1000 | #14-9965-82 | Invitrogen – Thermo |
| Rabbit IgG anti-CDHR3 polyclonal antibody | 1:1000 | #HPA011218 | Atlas - Sigma-Aldrich |
| Goat IgG anti-calnexin polyclonal antibody | 1:1000 | #PA5-19169 | Thermo |
| Rabbit IgG anti-Lamp1 (D3D11) monoclonal antibody | 1:1000 | #9091 | Cell Signaling |
| Mouse IgG2b anti-LC3b monoclonal antibody | 1:1000 | #83506 | Cell Signaling |
| Goat IgG anti-calnexin polyclonal antibody | 1:1000 | #PA5-19169 | Thermo |
| Mouse IgG2b anti-claudin-1 monoclonal antibody | 1:1000 | #sc-166338 | Santa Cruz |
| Rabbit IgG anti-ZO-1 polyclonal antibody | 1:200 | #61-7300 | Invitrogen |
| Mouse anti-β-Actin (clone AC-15) monoclonal antibody, peroxidase-conjugated | 1:15000 | #A3854 | Sigma-Aldrich |
| Rabbit IgG anti-TM173/STING monoclonal antibody | 1:1000 | #66680-1-Ig | Proteintech |
| Secondary Antibody | Dilution | Catalog # | Source |
| Recombinant mouse IgGk light chain binding protein conjugated to Horseradish Peroxidase (HRP) | 1:10000 | #sc516102 | Santa Cruz |
| Donkey anti-Goat IgG HRP- conjugated | 1:10000 | #A15999 | Thermo |
| Goat anti-rabbit IgG HRP- conjugated | 1:10000 | #32460 | Thermo |

### 9. Lactate dehydrogenase (LDH) release cytotoxicity assay

Apical and basolateral samples were used to quantify LDH release in RV-C15-infected HAE with the CytoTox 96® Non-Radioactive Cytotoxicity Assay kit (#G1780; Promega, Madison, WI) following the manufacturer’s protocol.

### 10. Transepithelial Electrical Resistance (TEER) assay

TEER was quantified in HAE using the Millicell Electrical Resistance System (ERS)-2 (Sigma) after adding 100µl of Pneumacult-ALI medium to the apical surface and incubating cultures for 30 minutes at 34°C. Statistical analysis of resulting data was done by ordinary one-way ANOVA and Tukey’s multiple comparison test methods using Prism GraphPad v.9 software (GraphPad).

### 11. Quantification of ZO-1 disposition using JAnaP

Junction coverage and characterization were quantified using the Junction Analyzer Program (JAnaP) as previously described [44–46]. In short, the perimeter of each cell was identified via waypoints in immunofluorescent images of ZO-1. The junctions were isolated from the background using a threshold value of 5-8. Threshold identification is described in the supplement of [44] and in the JAnaP User-Guide available at https://github.com/StrokaLab/JAnaP along with the JAnaP program in its entirety. Junction characterization was performed by calculating the length of each individual junction piece that coincides with the perimeter as well as the relative aspect ratio (RAR) with respect to the cell perimeter. A junction was classified as continuous if its length was greater than 15 pixels, otherwise it was deemed discontinuous and further separated into perpendicular or punctate if the RAR was greater or less than 1.2, respectively. Statistical analysis was done by one-way ANOVA and Dunnett’s multiple comparison test using Prism GraphPad v.9 software (GraphPad).

### 12. Measurement of Mucociliary Clearance (MCC)

Mucociliary transport was measured based on the transport of 2µm red-fluorescent polystyrene microspheres (Sigma-Aldrich). Five microliters of microspheres suspension (1:500 dilution in PBS; Sigma-Aldrich) was added on top of the native mucus; after 24-hour incubation at 34°C, videos of three regions were recorded using Zeiss Axio Observer 3 Inverted fluorescence microscope (Cell Observer HS image system, Zeiss Axiocam 503 mono camera, optimal acquisition mode, Zen 2007 software) equipped with 10x/0.25NA Ph1 air objective. Images were collected at a frame rate of 0.5 Hz for 10 seconds on the plane of the mucus gel. Images were acquired centrally within cultures and away from the edges, where mucus tends to accumulate. The microsphere tracking data analysis was based on an image processing algorithm that was custom written in MATLAB (The MathWorks). Briefly, the analysis software computes the XY-plane trajectories of each fluorescent microsphere in each frame. Using the trajectory data, displacement of microspheres was computed, and transport rate was calculated by dividing the displacement of microsphere by total time elapsed.

## Acknowledgments

We thank Drs. James Gern, Yuri A. Bochkov, Wai-Ming Lee, and Ann Palmenberg from University of Wisconsin – Madison (Madison, WI) for providing the rhinovirus plasmids. We would also like to thank the Addgene depositors Drs. Feng Zhang, Bob Weinberg, and Didier Trono for their contributions in making reagents broadly accessible and Drs. Matthew Walters and Ronald Crystal (Weill Cornell Medical College) for donating the BCi-NS1.1 cells. We are also grateful to the directors and teams at the Imaging Core, MPRI Flow Cytometry and Cell Sorting Facility, and the Laboratory for Biological Ultrastructure at the University of Maryland, College Park, for their assistance. We thank Dr. Michelle Itano (University of North Carolina, Chapel Hill) for helpful discussion regarding colocalization analysis. Finally, we acknowledge Eva Agostino for critical reading of the manuscript.

## Author contributions

TBG contributed to conceptualization and design of the study, as well as execution and analysis of the experiments. MEG developed HAE cultures with modified STING expression and analyzed rhinovirus infection in these cultures. DS performed and analyzed MCC experiments. KMG and JWJ evaluated ZO-1 disposition using the JANaP software. KMS validated JANaP results and contributed expertise. GAD validated MCC data and contributed expertise. MAS contributed to the study design and obtained funding. TBG and MAS wrote the manuscript. All authors reviewed and approved the final manuscript.

## Supplementary files

### I. Supplementary tables

**S1 Table.**
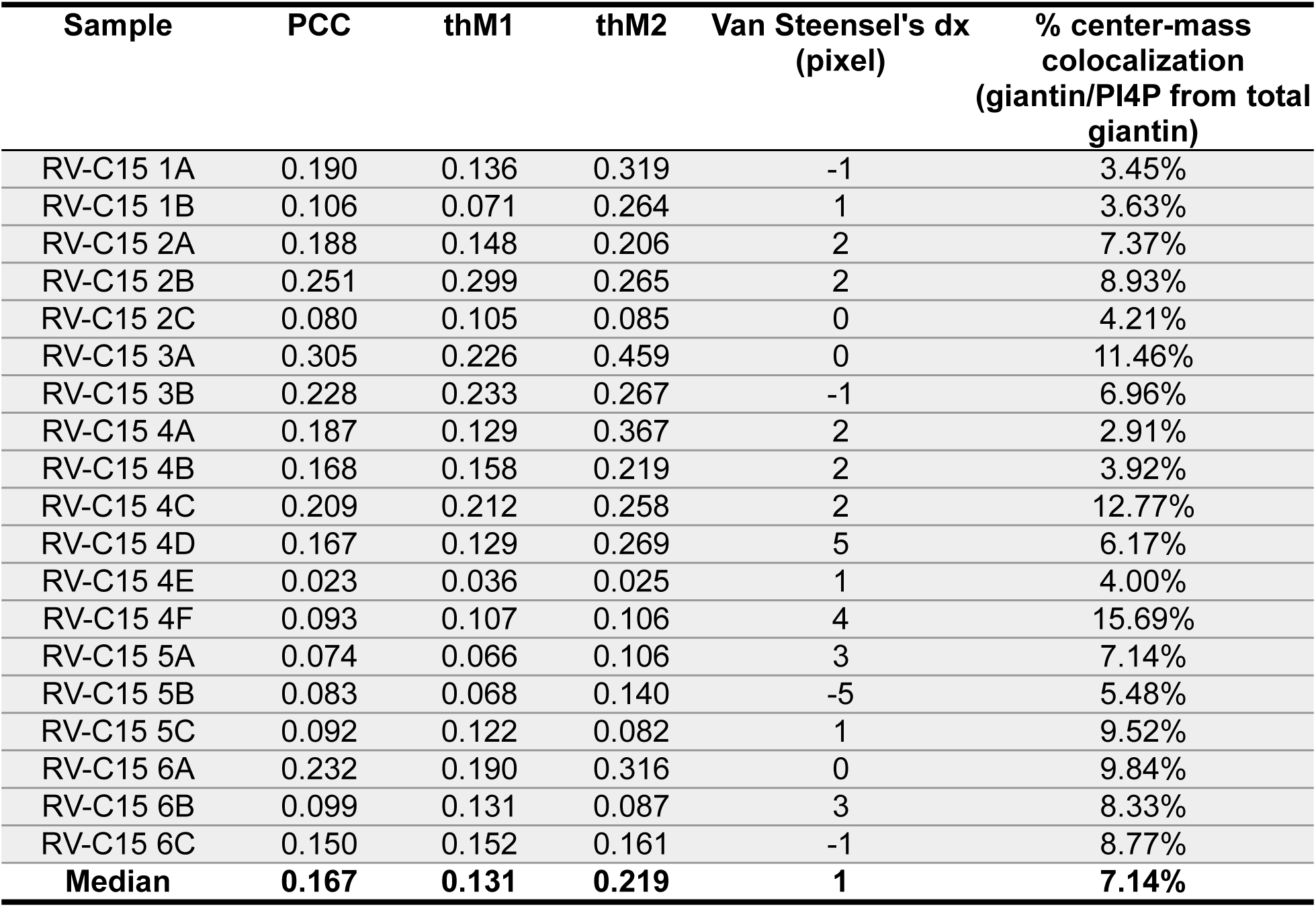
Pixel-intensity based and spatial (distance between center-mass) colocalization analysis between giantin and PI4P in RV-C15 infected HAE.

**S2 Table.** Pixel-intensity based and spatial (distance between center-mass) colocalization analysis between giantin and PI4P in RV-A16 infected HAE.

| Sample | PCC | thM1 | thM2 | Van Steensel's dx<br>(pixel) | % center-mass<br>colocalization<br>(giantin/PI4P from<br>total giantin) |
| --- | --- | --- | --- | --- | --- |
| RV-A16 1A | 0.053 | 0.066 | 0.068 | -15 | 3.45% |
| RV-A16 1B | 0.133 | 0.083 | 0.266 | -4 | 2.50% |
| RV-A16 1C | 0.081 | 0.073 | 0.135 | 2 | 9.43% |
| RV-A16 2A | 0.051 | 0.076 | 0.093 | -17 | 3.75% |
| RV-A16 2B | 0.102 | 0.095 | 0.150 | 9 | 0.00% |
| RV-A16 2C | 0.062 | 0.049 | 0.109 | 1 | 2.22% |
| RV-A16 3A | 0.206 | 0.166 | 0.328 | -2 | 7.69% |
| RV-A16 3B | 0.035 | 0.028 | 0.110 | 17 | 0.00% |
| RV-A16 3C | 0.210 | 0.133 | 0.421 | 3 | 6.25% |
| RV-A16 3D | 0.192 | 0.151 | 0.271 | 1 | 0.00% |
| RV-A16 4A | 0.086 | 0.063 | 0.189 | 20 | 3.17% |
| RV-A16 4B | 0.081 | 0.046 | 0.187 | 6 | 2.60% |
| RV-A16 4C | 0.122 | 0.054 | 0.319 | -2 | 4.11% |
| **Median** | **0.086** | **0.073** | **0.187** | **1** | **3.17%** |

**S3 Table.** Pixel-intensity based and spatial (distance between center-mass) colocalization analysis between giantin and PI4P in RV-A2 infected HAE.

| Sample | PCC | thM1 | thM2 | Van Steensel's dx<br>(pixel) | % center-mass<br>colocalization<br>(giantin/PI4P from<br>total giantin) |
| --- | --- | --- | --- | --- | --- |
| RV-A2 1A | 0.141 | 0.165 | 0.136 | 2 | 16.67% |
| RV-A2 1B | 0.116 | 0.125 | 0.122 | 4 | 6.00% |
| RV-A2 2A | 0.066 | 0.059 | 0.113 | 2 | 2.19% |
| RV-A2 2B | 0.206 | 0.216 | 0.231 | -3 | 2.50% |
| RV-A2 3A | 0.039 | 0.056 | 0.035 | 1 | 3.85% |
| RV-A2 4A | 0.028 | 0.032 | 0.041 | 5 | 3.70% |
| RV-A2 5A | 0.095 | 0.092 | 0.122 | -1 | 7.14% |
| RV-A2 6A | 0.180 | 0.279 | 0.158 | 0 | 5.26% |
| **Median** | **0.106** | **0.109** | **0.122** | **2** | **4.55%** |

**S4 Table.** Pixel-intensity based and spatial (distance between center-mass) colocalization analysis between dsRNA and giantin in RV-C15 infected HAE.

| **Sample** | **PCC** | **thM1** | **thM2** | **Van Steensel's dx**<br>**(pixel)** | **% center-mass**<br>**colocalization**<br>**(dsRNA/giantin from**<br>**total dsRNA)** |
| --- | --- | --- | --- | --- | --- |
| RV-C15 1A | 0.174 | 0.157 | 0.228 | -1 | 3.65% |
| RV-C15 1B | 0.175 | 0.153 | 0.248 | 1 | 11.39% |
| RV-C15 2A | 0.168 | 0.158 | 0.227 | 2 | 13.04% |
| RV-C15 2B | 0.237 | 0.354 | 0.191 | 2 | 11.83% |
| RV-C15 2C | 0.176 | 0.223 | 0.158 | 0 | 23.53% |
| RV-C15 3A | 0.117 | 0.120 | 0.149 | -1 | 14.85% |
| RV-C15 3B | 0.133 | 0.167 | 0.142 | 2 | 5.62% |
| RV-C15 4A | 0.094 | 0.125 | 0.102 | 1 | 8.43% |
| RV-C15 4B | 0.142 | 0.122 | 0.215 | 1 | 11.11% |
| RV-C15 4C | 0.150 | 0.165 | 0.187 | 8 | 12.90% |
| RV-C15 4D | 0.106 | 0.124 | 0.117 | 1 | 5.36% |
| RV-C15 4E | 0.105 | 0.080 | 0.168 | 0 | 2.02% |
| RV-C15 4F | 0.056 | 0.069 | 0.071 | 2 | 11.46% |
| RV-C15 5A | 0.062 | 0.044 | 0.128 | -3 | 8.54% |
| RV-C15 5B | 0.079 | 0.071 | 0.121 | -7 | 2.47% |
| RV-C15 5C | 0.102 | 0.083 | 0.162 | -1 | 4.81% |
| RV-C15 6A | 0.162 | 0.163 | 0.185 | 0 | 6.93% |
| RV-C15 6B | 0.060 | 0.056 | 0.094 | 3 | 6.59% |
| RV-C15 6C | 0.085 | 0.056 | 0.163 | 1 | 10.87% |
| **Median** | **0.117** | **0.124** | **0.162** | **1** | **8.54%** |

**S5 Table.** Pixel-intensity based and spatial (distance between center-mass) colocalization analysis between dsRNA and giantin in RV-A16 infected HAE.

| Sample | PCC | thM1 | thM2 | Van Steensel's dx<br>(pixel) | % center-mass<br>colocalization<br>(dsRNA/giantin from<br>total dsRNA) |
| --- | --- | --- | --- | --- | --- |
| RV-A16 1A | 0.069 | 0.056 | 0.146 | -12 | 6.67% |
| RV-A16 1B | 0.133 | 0.092 | 0.235 | 0 | 5.47% |
| RV-A16 1C | 0.069 | 0.055 | 0.147 | 2 | 3.96% |
| RV-A16 2A | 0.140 | 0.128 | 0.241 | 0 | 3.41% |
| RV-A16 2B | 0.050 | 0.049 | 0.104 | 1 | 4.65% |
| RV-A16 2C | 0.032 | 0.023 | 0.097 | 0 | 5.08% |
| RV-A16 3A | 0.161 | 0.173 | 0.194 | -14 | 10.20% |
| RV-A16 3B | 0.138 | 0.079 | 0.308 | 1 | 15.38% |
| RV-A16 3C | 0.007 | 0.024 | 0.043 | 20 | 2.04% |
| RV-A16 3D | 0.060 | 0.056 | 0.091 | -9 | 5.13% |
| RV-A16 4A | 0.105 | 0.122 | 0.116 | 4 | 6.48% |
| RV-A16 4B | 0.004 | 0.010 | 0.018 | 15 | 2.22% |
| RV-A16 4C | 0.074 | 0.045 | 0.149 | -19 | 2.94% |
| **Median** | **0.069** | **0.056** | **0.146** | **0** | **5.08%** |

**S6 Table.** Pixel-intensity based and spatial (distance between center-mass) colocalization analysis between dsRNA and giantin in RV-A2 infected HAE.

| Sample | PCC | thM1 | thM2 | Van Steensel's dx<br>(pixel) | % center-mass<br>colocalization<br>(dsRNA/giantin from<br>total dsRNA) |
| --- | --- | --- | --- | --- | --- |
| RV-A2 1A | 0.032 | 0.036 | 0.058 | 2 | 3.54% |
| RV-A2 1B | 0.045 | 0.038 | 0.088 | 14 | 4.26% |
| RV-A2 2A | 0.053 | 0.058 | 0.077 | 1 | 11.54% |
| RV-A2 2B | 0.046 | 0.074 | 0.054 | -19 | 5.15% |
| RV-A2 3A | 0.044 | 0.029 | 0.108 | 1 | 1.30% |
| RV-A2 4A | 0.087 | 0.066 | 0.140 | 0 | 8.62% |
| RV-A2 5A | 0.075 | 0.046 | 0.190 | 19 | 7.50% |
| RV-A2 6A | 0.106 | 0.179 | 0.110 | 10 | 5.21% |
| **Median** | **0.050** | **0.052** | **0.098** | **2** | **5.18%** |

**S7 Table.** Pixel-intensity based and spatial (distance between center-mass) colocalization analysis between dsRNA and PI4P in RV-C15 infected HAE.

| **Sample** | **PCC** | **thM1** | **thM2** | **Van Steensel's dx**<br>**(pixel)** | **% center-mass**<br>**colocalization**<br>**(dsRNA/PI4P from**<br>**total dsRNA)** |
| --- | --- | --- | --- | --- | --- |
| RV-C15 1A | 0.259 | 0.357 | 0.220 | 0 | 4.38% |
| RV-C15 1B | 0.108 | 0.224 | 0.098 | -2 | 3.80% |
| RV-C15 2A | 0.281 | 0.371 | 0.256 | 0 | 10.14% |
| RV-C15 2B | 0.280 | 0.385 | 0.235 | -1 | 4.30% |
| RV-C15 2C | 0.051 | 0.068 | 0.060 | -1 | 9.80% |
| RV-C15 3A | 0.212 | 0.298 | 0.182 | 0 | 7.92% |
| RV-C15 3B | 0.230 | 0.290 | 0.215 | -1 | 4.49% |
| RV-C15 4A | 0.253 | 0.519 | 0.148 | 0 | 19.28% |
| RV-C15 4B | 0.227 | 0.220 | 0.280 | -1 | 11.11% |
| RV-C15 4C | 0.292 | 0.327 | 0.305 | -1 | 19.35% |
| RV-C15 4D | 0.244 | 0.388 | 0.176 | 0 | 2.54% |
| RV-C15 4E | 0.040 | 0.029 | 0.087 | -2 | 2.02% |
| RV-C15 4F | 0.052 | 0.065 | 0.067 | 0 | 3.13% |
| RV-C15 5A | 0.211 | 0.168 | 0.302 | -2 | 7.32% |
| RV-C15 5B | 0.092 | 0.122 | 0.101 | -1 | 2.47% |
| RV-C15 5C | 0.082 | 0.055 | 0.161 | -1 | 3.85% |
| RV-C15 6A | 0.295 | 0.371 | 0.254 | 0 | 7.92% |
| RV-C15 6B | 0.144 | 0.097 | 0.243 | 0 | 8.79% |
| RV-C15 6C | 0.130 | 0.085 | 0.233 | 6 | 10.87% |
| **Median** | **0.212** | **0.224** | **0.215** | **-1** | **7.32%** |

**S8 Table.** Pixel-intensity based and spatial (distance between center-mass) colocalization analysis between dsRNA and PI4P in RV-A16 infected HAE.

| Sample | PCC | thM1 | thM2 | Van Steensel's dx<br>(pixel) | % center-mass<br>colocalization<br>(dsRNA/PI4P from<br>total dsRNA) |
| --- | --- | --- | --- | --- | --- |
| RV-A16 1A | 0.068 | 0.056 | 0.142 | -1 | 6.67% |
| RV-A16 1B | 0.105 | 0.146 | 0.117 | 1 | 3.97% |
| RV-A16 1C | 0.194 | 0.184 | 0.262 | -1 | 2.97% |
| RV-A16 2A | 0.095 | 0.111 | 0.170 | -1 | 0.00% |
| RV-A16 2B | 0.099 | 0.108 | 0.144 | 0 | 4.65% |
| RV-A16 2C | 0.048 | 0.052 | 0.098 | 0 | 3.39% |
| RV-A16 3A | 0.243 | 0.359 | 0.204 | 2 | 6.12% |
| RV-A16 3B | 0.116 | 0.154 | 0.154 | -13 | 7.69% |
| RV-A16 3C | 0.184 | 0.296 | 0.165 | 0 | 2.04% |
| RV-A16 3D | 0.193 | 0.218 | 0.195 | -1 | 6.41% |
| RV-A16 4A | 0.196 | 0.382 | 0.122 | 0 | 2.78% |
| RV-A16 4B | 0.112 | 0.192 | 0.084 | 0 | 2.22% |
| RV-A16 4C | 0.201 | 0.290 | 0.163 | 2 | 5.88% |
| **Median** | **0.116** | **0.184** | **0.154** | **0** | **3.97%** |

**S9 Table.** Pixel-intensity based and spatial (distance between center-mass) colocalization analysis between dsRNA and PI4P in RV-A2 infected HAE.

| Sample | PCC | thM1 | thM2 | Van Steensel's dx<br>(pixel) | % center-mass<br>colocalization<br>(dsRNA/PI4P from<br>total dsRNA) |
| --- | --- | --- | --- | --- | --- |
| RV-A2 1A | 0.072 | 0.060 | 0.118 | -1 | 1.77% |
| RV-A2 1B | 0.077 | 0.058 | 0.137 | -2 | 3.19% |
| RV-A2 2A | 0.110 | 0.152 | 0.106 | -1 | 12.82% |
| RV-A2 2B | 0.118 | 0.163 | 0.111 | -1 | 6.19% |
| RV-A2 3A | 0.049 | 0.024 | 0.141 | -1 | 2.60% |
| RV-A2 4A | 0.079 | 0.069 | 0.115 | -2 | 4.31% |
| RV-A2 5A | 0.081 | 0.058 | 0.182 | 0 | 5.63% |
| RV-A2 6A | 0.211 | 0.224 | 0.243 | -1 | 16.13% |
| **Median** | **0.080** | **0.065** | **0.128** | **-1** | **4.97%** |

**S10 Table.**
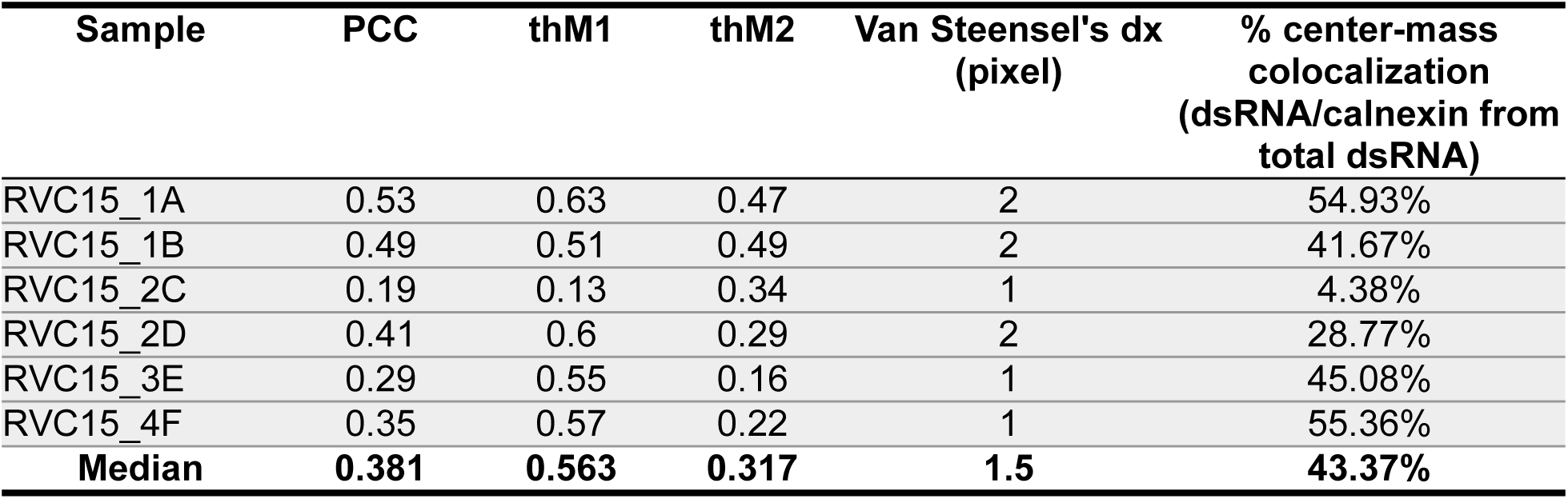
Pixel-intensity based and spatial (distance between center-mass) colocalization analysis between dsRNA and calnexin in RV-C15-infected HAE.

**S11 Table.** Pixel-intensity based and spatial (distance between center-mass) colocalization analysis between dsRNA and calnexin in RV-A16-infected HAE.

| Sample | PCC | thM1 | thM2 | Van Steensel's dx<br>(pixel) | % center-mass<br>colocalization<br>(dsRNA/calnexin from<br>total dsRNA) |
| --- | --- | --- | --- | --- | --- |
| RV-A16_1A | 0.366 | 0.407 | 0.387 | 1 | 14.18% |
| RV-A16_1B | 0.289 | 0.372 | 0.292 | 3 | 10.16% |
| RV-A16_1C | 0.265 | 0.328 | 0.271 | 2 | 2.97% |
| RV-A16_1D | 0.340 | 0.231 | 0.607 | 3 | 2.68% |
| RV-A16_1E | 0.287 | 0.276 | 0.374 | 2 | 3.26% |
| RV-A16_2F | 0.352 | 0.263 | 0.618 | 2 | 3.15% |
| RV-A16_2G | 0.320 | 0.236 | 0.584 | 4 | 0.83% |
| RV-A16_2H | 0.358 | 0.255 | 0.637 | 3 | 0.55% |
| RV-A16_3I | 0.339 | 0.389 | 0.359 | 2 | 22.35% |
| RV-A16_3J | 0.272 | 0.315 | 0.312 | 2 | 1.05% |
| RV-A16_3K | 0.364 | 0.280 | 0.633 | 0 | 4.04% |
| **Median** | **0.339** | **0.280** | **0.387** | **2** | **3.15%** |

**S12 Table.**
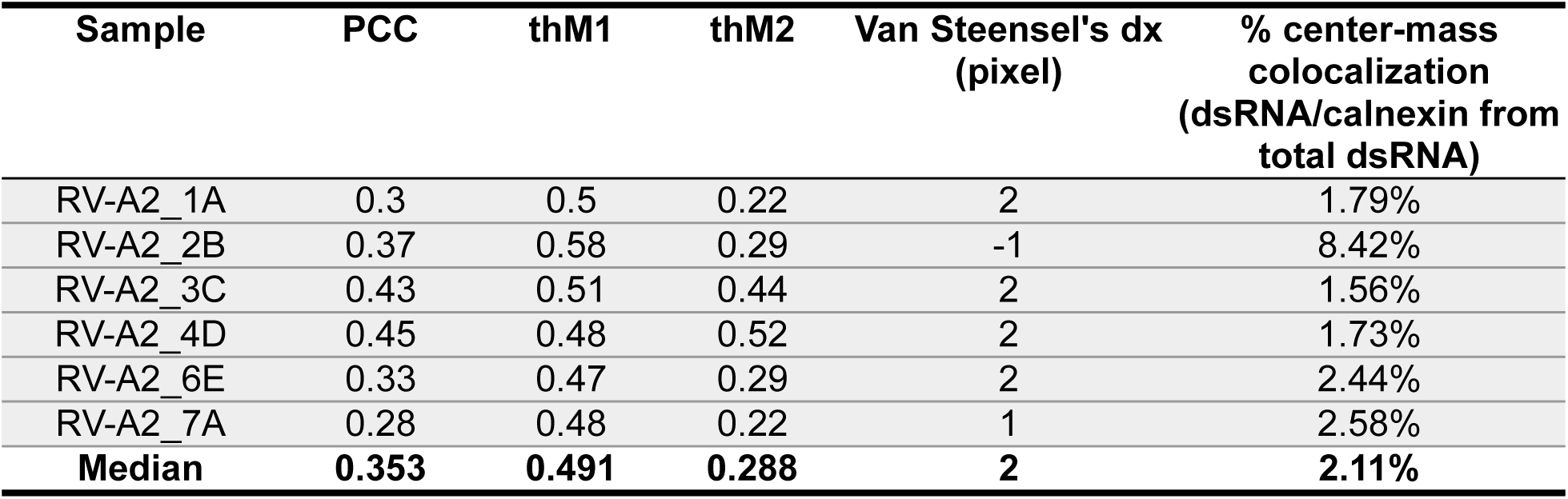
Pixel-intensity based and spatial (distance between center-mass) colocalization analysis between dsRNA and calnexin in RV-A2-infected HAE.

| Sample | PCC | thM1 | thM2 | Van Steensel's dx<br>(pixel) | % center-mass<br>colocalization<br>(dsRNA/calnexin from<br>total dsRNA) |
| --- | --- | --- | --- | --- | --- |
| RV-A2_1A | 0.3 | 0.5 | 0.22 | 2 | 1.79% |
| RV-A2_2B | 0.37 | 0.58 | 0.29 | -1 | 8.42% |
| RV-A2_3C | 0.43 | 0.51 | 0.44 | 2 | 1.56% |
| RV-A2_4D | 0.45 | 0.48 | 0.52 | 2 | 1.73% |
| RV-A2_6E | 0.33 | 0.47 | 0.29 | 2 | 2.44% |
| RV-A2_7A | 0.28 | 0.48 | 0.22 | 1 | 2.58% |
| **Median** | **0.353** | **0.491** | **0.288** | **2** | **2.11%** |

**S13 Table.** Pixel-intensity based and spatial (distance between center-mass) colocalization analysis between dsRNA and giantin in RV-C15- infected HAE.

| Sample | PCC | thM1 | thM2 | Van Steensel's dx<br>(pixel) | % center-mass<br>colocalization<br>(dsRNA/giantin from<br>total dsRNA) |
| --- | --- | --- | --- | --- | --- |
| RVC15_1A | 0.14 | 0.15 | 0.15 | -1 | 18.31% |
| RVC15_1B | 0.19 | 0.2 | 0.2 | -3 | 12.50% |
| RVC15_2C | 0.07 | 0.06 | 0.15 | 1 | 2.51% |
| RVC15_2D | 0.13 | 0.2 | 0.1 | 0 | 10.96% |
| RVC15_3E | 0.09 | 0.17 | 0.06 | -2 | 24.59% |
| RVC15_4F | 0.15 | 0.23 | 0.11 | -1 | 25.23% |
| **Median** | **0.136** | **0.183** | **0.130** | **-1** | **15.40%** |

**S14 Table.** Pixel-intensity based and spatial (distance between center-mass) colocalization analysis between dsRNA and giantin in RV-A16-infected HAE.

| Sample | PCC | thM1 | thM2 | Van Steensel's dx<br>(pixel) | % center-mass<br>colocalization<br>(dsRNA/giantin from<br>total dsRNA) |
| --- | --- | --- | --- | --- | --- |
| RV-A16_1A | 0.372 | 0.501 | 0.333 | -3 | 10.64% |
| RV-A16_1B | 0.331 | 0.441 | 0.313 | 2 | 1.56% |
| RV-A16_1C | 0.363 | 0.607 | 0.264 | -1 | 6.93% |
| RV-A16_1D | 0.241 | 0.306 | 0.311 | 1 | 0.24% |
| RV-A16_1E | 0.269 | 0.480 | 0.223 | -1 | 14.67% |
| RV-A16_2F | 0.319 | 0.346 | 0.450 | -1 | 2.45% |
| RV-A16_2G | 0.376 | 0.378 | 0.515 | 5 | 3.32% |
| RV-A16_2H | 0.241 | 0.318 | 0.337 | 0 | 0.00% |
| RV-A16_3I | 0.115 | 0.172 | 0.153 | 0 | 8.24% |
| RV-A16_3J | 0.215 | 0.311 | 0.228 | 5 | 21.05% |
| RV-A16_3K | 0.259 | 0.260 | 0.440 | -18 | 1.01% |
| **Median** | **0.269** | **0.346** | **0.313** | **0** | **3.32%** |

**S15 Table.** Pixel-intensity based and spatial (distance between center-mass) colocalization analysis between dsRNA and giantin in RV-A2- infected HAE.

| Sample | PCC | thM1 | thM2 | Van Steensel's dx<br>(pixel) | % center-mass<br>colocalization<br>(dsRNA/giantin from<br>total dsRNA) |
| --- | --- | --- | --- | --- | --- |
| RV-A2_1A | 0.19 | 0.27 | 0.21 | 0 | 4.11% |
| RV-A2_2B | 0.33 | 0.52 | 0.27 | -3 | 2.11% |
| RV-A2_3C | 0.27 | 0.3 | 0.33 | 2 | 1.56% |
| RV-A2_4D | 0.34 | 0.36 | 0.44 | 0 | 8.67% |
| RV-A2_6E | 0.26 | 0.39 | 0.23 | -1 | 7.32% |
| RV-A2_7A | 0.22 | 0.39 | 0.18 | 1 | 2.58% |
| **Median** | **0.266** | **0.378** | **0.249** | **0** | **3.35%** |

**S16 Table.** Pixel-intensity based and spatial (distance between center-mass) colocalization analysis between dsRNA and STING in RV-C15-infected HAE.

| **Sample** | **PCC** | **thM1** | **thM2** | **Van Steensel's dx**<br>**(pixel)** | **% center-mass**<br>**colocalization**<br>**(dsRNA/STING from**<br>**total dsRNA)** |
| --- | --- | --- | --- | --- | --- |
| RVC15_1A | 0.241 | 0.256 | 0.288 | 0 | 3.74% |
| RVC15_1B | 0.363 | 0.492 | 0.326 | 2 | 0.92% |
| RVC15_1C | 0.297 | 0.207 | 0.597 | -4 | 11.25% |
| RVC15_1D | 0.025 | 0.029 | 0.054 | 2 | 6.00% |
| RVC15_1E | 0.055 | 0.055 | 0.090 | 3 | 3.68% |
| RVC15_1F | 0.041 | 0.044 | 0.087 | 0 | 2.97% |
| RVC15_1G | 0.060 | 0.030 | 0.163 | -2 | 1.32% |
| RVC15_1H | 0.005 | 0.023 | 0.021 | -1 | 1.46% |
| RVC15_2A | 0.321 | 0.406 | 0.319 | -1 | 7.55% |
| RVC15_2B | 0.367 | 0.395 | 0.455 | -1 | 6.56% |
| RVC15_2C | 0.239 | 0.197 | 0.333 | 2 | 2.90% |
| RVC15_2D | 0.212 | 0.232 | 0.300 | -7 | 3.70% |
| RVC15_2E | 0.005 | 0.021 | 0.049 | -3 | 5.17% |
| RVC15_2F | 0.127 | 0.096 | 0.213 | -1 | 4.65% |
| RVC15_3A | 0.402 | 0.471 | 0.400 | 2 | 4.76% |
| RVC15_3B | 0.314 | 0.268 | 0.466 | -3 | 2.04% |
| RVC15_3C | 0.265 | 0.266 | 0.325 | 0 | 3.28% |
| RVC15_3D | 0.198 | 0.152 | 0.300 | 1 | 4.94% |
| RVC15_3E | 0.455 | 0.631 | 0.371 | 1 | 0.00% |
| RVC15_3F | 0.149 | 0.292 | 0.101 | 1 | 4.29% |
| **Median** | **0.226** | **0.220** | **0.300** | **0** | **3.72%** |

**S17 Table.** Pixel-intensity based and spatial (distance between center-mass) colocalization analysis between Lamp1 and LC3b in RV-C15-infected HAE.

| **Sample** | **PCC** | **thM1** | **thM2** | **Van Steensel's dx**<br>**(pixel)** | **% center-mass**<br>**colocalization**<br>**(Lamp1/LC3b from**<br>**total Lamp1)** |
| --- | --- | --- | --- | --- | --- |
| RV-C15-1A | 0.262 | 0.129 | 0.570 | -1 | 5.17% |
| RV-C15-1B | 0.040 | 0.032 | 0.098 | -1 | 1.43% |
| RV-C15-1C | 0.159 | 0.114 | 0.300 | 0 | 3.57% |
| RV-C15-1D | 0.048 | 0.016 | 0.240 | 0 | 0.00% |
| RV-C15-2A | 0.058 | 0.034 | 0.183 | -2 | 4.55% |
| RV-C15-2B | 0.256 | 0.120 | 0.631 | -1 | 16.67% |
| RV-C15-2C | 0.027 | 0.016 | 0.107 | -1 | 0.00% |
| RV-C15-3A | 0.148 | 0.037 | 0.684 | -1 | 0.00% |
| RV-C15-3B | 0.157 | 0.047 | 0.620 | -1 | 3.06% |
| RV-C15-3C | 0.107 | 0.023 | 0.596 | -2 | 12.50% |
| RV-C15-3D | 0.186 | 0.071 | 0.577 | -2 | 3.13% |
| RV-C15-4A | 0.128 | 0.030 | 0.677 | -2 | 2.50% |
| RV-C15-4B | 0.142 | 0.040 | 0.597 | -2 | 0.00% |
| RV-C15-4C | 0.186 | 0.061 | 0.690 | -3 | 2.50% |
| RV-C15-5A | 0.249 | 0.082 | 0.792 | -1 | 2.60% |
| RV-C15-5B | 0.067 | 0.010 | 0.534 | -1 | 0.00% |
| RV-C15-6A | 0.321 | 0.153 | 0.723 | -1 | 3.13% |
| RV-C15-6B | 0.054 | 0.012 | 0.323 | 0 | 0.00% |
| RV-C15-6C | 0.024 | 0.007 | 0.129 | -1 | 0.00% |
| **Median** | **0.142** | **0.037** | **0.577** | **-1** | **2.50%** |

**S18 Table.** Pixel-intensity based and spatial (distance between center-mass) colocalization analysis between Lamp1 and LC3b in RV-A16-infected HAE.

| Sample | PCC | thM1 | thM2 | Van Steensel's dx<br>(pixel) | % center-mass<br>colocalization<br>(Lamp1/LC3b from<br>total Lamp1) |
| --- | --- | --- | --- | --- | --- |
| RV-A16-1A | 0.604 | 0.670 | 0.569 | -1 | 16.67% |
| RV-A16-1B | 0.388 | 0.585 | 0.296 | -1 | 9.38% |
| RV-A16-1C | 0.399 | 0.635 | 0.290 | -2 | 0.00% |
| RV-A16-3A | 0.510 | 0.610 | 0.488 | -1 | 24.14% |
| RV-A16-3B | 0.435 | 0.584 | 0.356 | -1 | 17.07% |
| RV-A16-3C | 0.409 | 0.557 | 0.338 | -2 | 8.33% |
| RV-A16-3D | 0.583 | 0.665 | 0.538 | -2 | 27.27% |
| RV-A16-4A | 0.568 | 0.647 | 0.530 | -1 | 16.67% |
| RV-A16-4C | 0.535 | 0.618 | 0.493 | -1 | 7.41% |
| RV-A16-5A | 0.395 | 0.590 | 0.308 | -2 | 13.73% |
| RV-A16-5B | 0.418 | 0.605 | 0.318 | -2 | 2.86% |
| RV-A16-5C | 0.392 | 0.599 | 0.304 | -2 | 17.14% |
| RV-A16-5D | 0.456 | 0.639 | 0.376 | -2 | 7.32% |
| **Median** | **0.435** | **0.610** | **0.356** | **-2** | **13.73%** |

**S19 Table.** Pixel-intensity based and spatial (distance between center-mass) colocalization analysis between Lamp 1 and LC3b in RV-A2-infected HAE.

| Sample | PCC | thM1 | thM2 | Van Steensel's dx<br>(pixel) | % center-mass<br>colocalization<br>(Lamp1/LC3b from<br>total Lamp1) |
| --- | --- | --- | --- | --- | --- |
| RV-A2-1A | 0.361 | 0.372 | 0.388 | 0 | 9.76% |
| RV-A2-2A | 0.370 | 0.175 | 0.869 | -1 | 2.13% |
| RV-A2-3A | 0.393 | 0.281 | 0.601 | -1 | 3.80% |
| RV-A2-3B | 0.228 | 0.225 | 0.299 | 0 | 0.00% |
| RV-A2-4A | 0.166 | 0.096 | 0.367 | -2 | 0.00% |
| RV-A2-4b | 0.141 | 0.119 | 0.231 | -2 | 0.00% |
| RV-A2-4C | 0.135 | 0.120 | 0.242 | -3 | 4.08% |
| **Median** | **0.228** | **0.175** | **0.367** | **-1** | **2.13%** |

### II. Supplementary figures

**S1 Figure.**
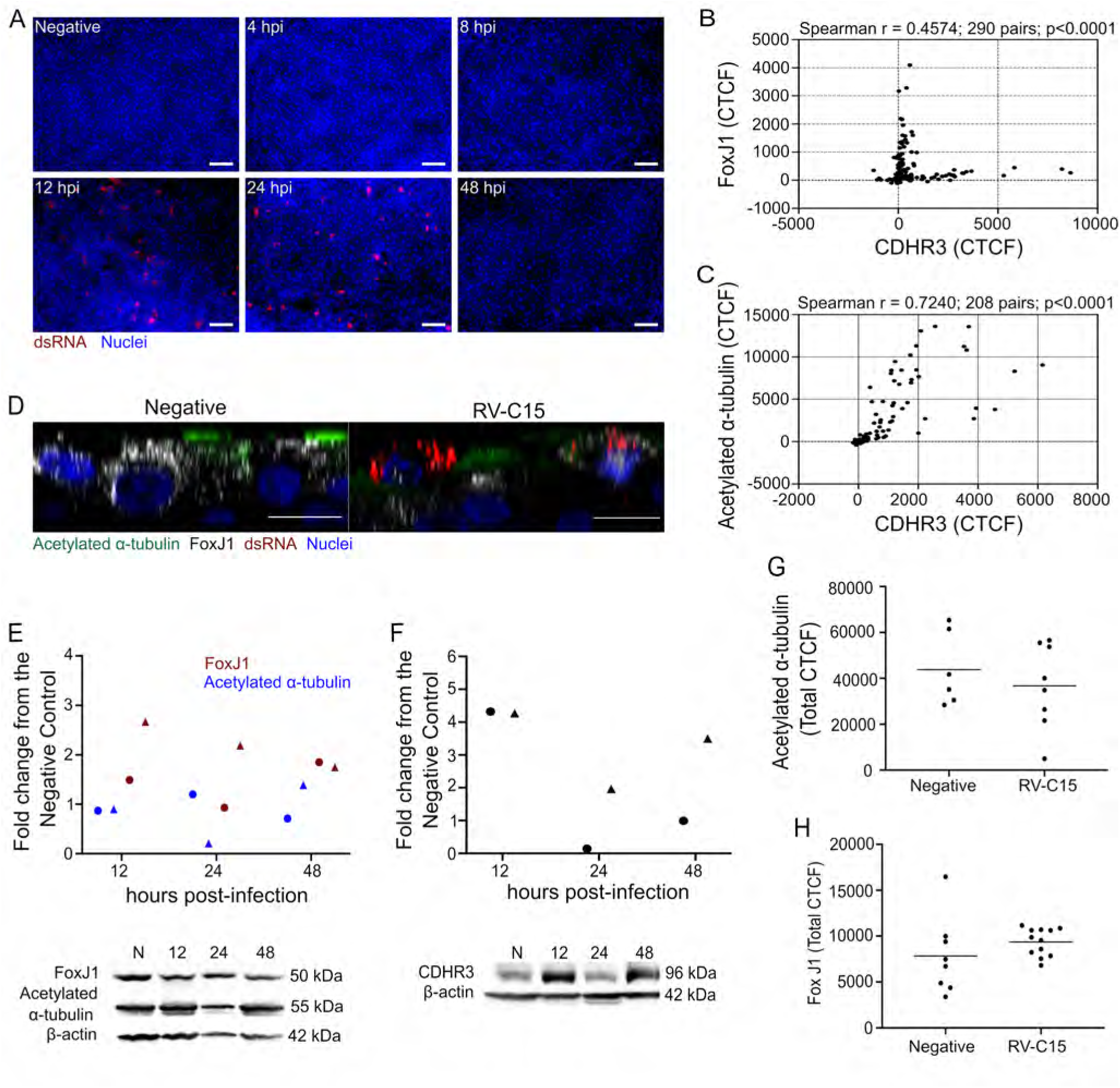
RV-C15 replicates in ciliated epithelial cells, leading to decreased CDHR3 levels. **A**: Immunofluorescence to detect dsRNA (red) in RV-C15 (10^10^ RNA) infected HAE (nuclei, blue; scale bar = 50µm). **B-C**: Spearman correlation analysis (two-tailed; 0.95% confidence interval) between fluorescence levels of CDHR3 and FoxJ1 (**B**) or acetylated α-tubulin (**C**) in non-infected HAE at 12 hpi. **D**: Orthogonal XY view from non-infected and RV-C15-infected (dsRNA+, red) HAE (nuclei, blue) stained by immunofluorescence for FoxJ1 (gray) and acetylated α-tubulin (green) at 12 hpi (z-stacks at 1µm of thickness; scale bar = 5µm). **E-F:** Fold change graph represents FoxJ1 (**E**), acetylated α-tubulin (**E**), and CDHR3 (**F**) protein levels normalized to the endogenous control (actin) and compared to non-infected cultures. Data shown are from two independent donors, represented by circles and triangles. Blot below is from the donor represented by circles (**E**) and triangles (**F**). **G-H**: Quantification of fluorescence levels (CTCF) for FoxJ1 (**G**) and acetylated α-tubulin (**H**) in non-infected and RV-C15-infected (dsRNA+) HAE at 12 hpi. Statistical analysis was done using Mann-Whitney U test (Two-tailed; 0.95% confidence interval).

**S2 Figure.**
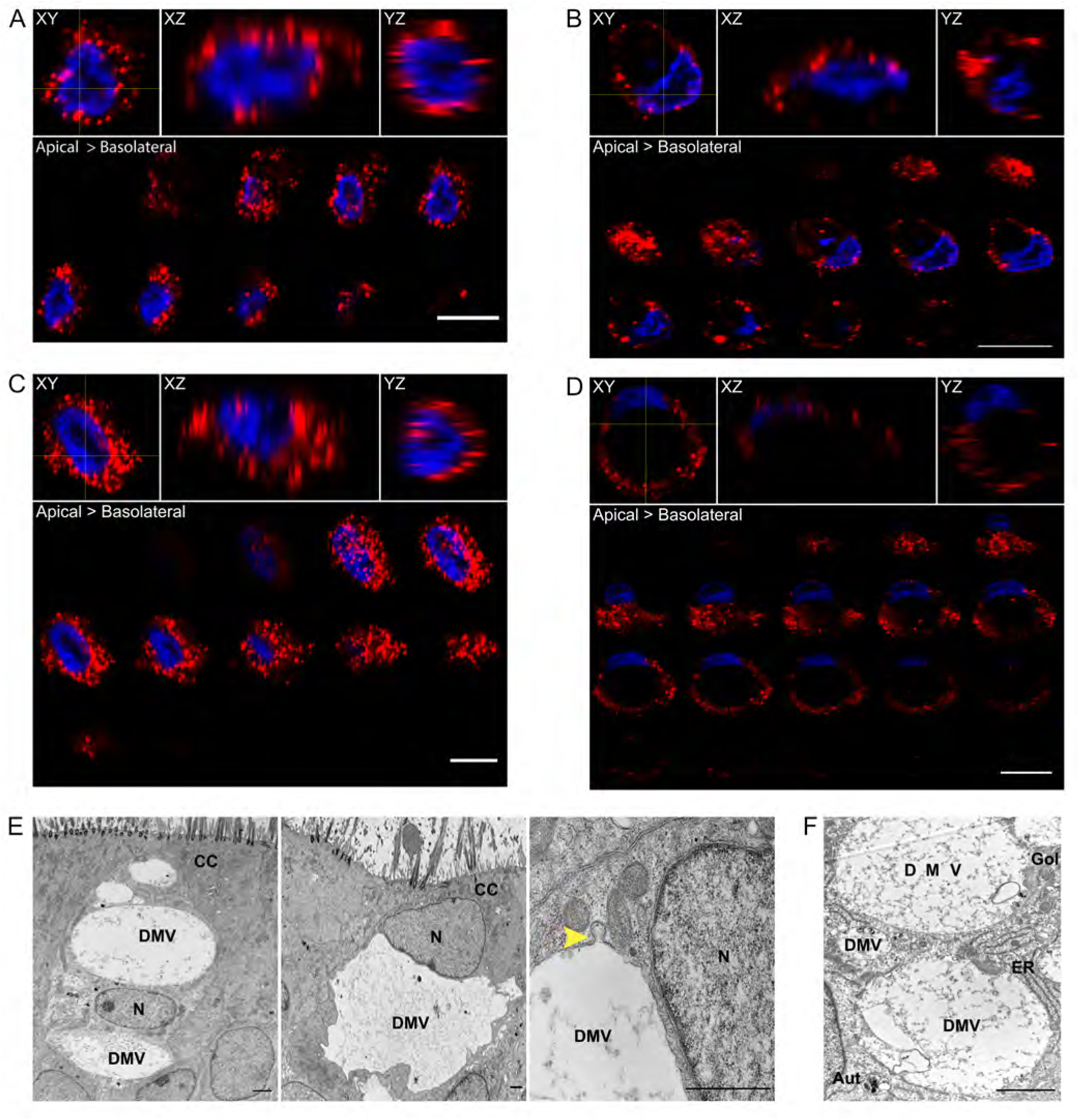
“Ring-like” dsRNA localization in HAE is not specific to RV-C15 infection. **A-D**: Orthogonal views (XY, XZ and YZ planes; yellow lines show the location of XZ and YZ views on the XY plane) from the RV-A16 **(A-B)** and RV-A2 (**C-D**) -infected HAE immunostained for dsRNA (red; nuclei, blue) at 12 hpi (z-stacks at 1µm of thickness). Z-stacks at 1µm of thickness shows two profiles for dsRNA (red) detection by immunofluorescence at perinuclear **(A, C**) or close to the plasma membrane in a “ring-like” disposition (**B, D**) at 12 hpi (scale bar = 10µm). **E-F:** Transmission electron microscopy of HAE infected with RV-A16 (**E**) or RV-A2 (**F**) at 12 hpi (scale bar = 5µm). Visualization of ciliated cells *(CC*) with large, double-membrane vesicles (DMV; **E and F**) located above (**E –** *left panel)* or below the nucleus (**E –** *middle panel*); and the fusion of small vesicles to the big ones (**E –** *right panel; yellow arrow).* N = nucleus; Gol = golgi; ER = endoplasmic reticulum; Aut = autophagosome.

**S3 Figure.**
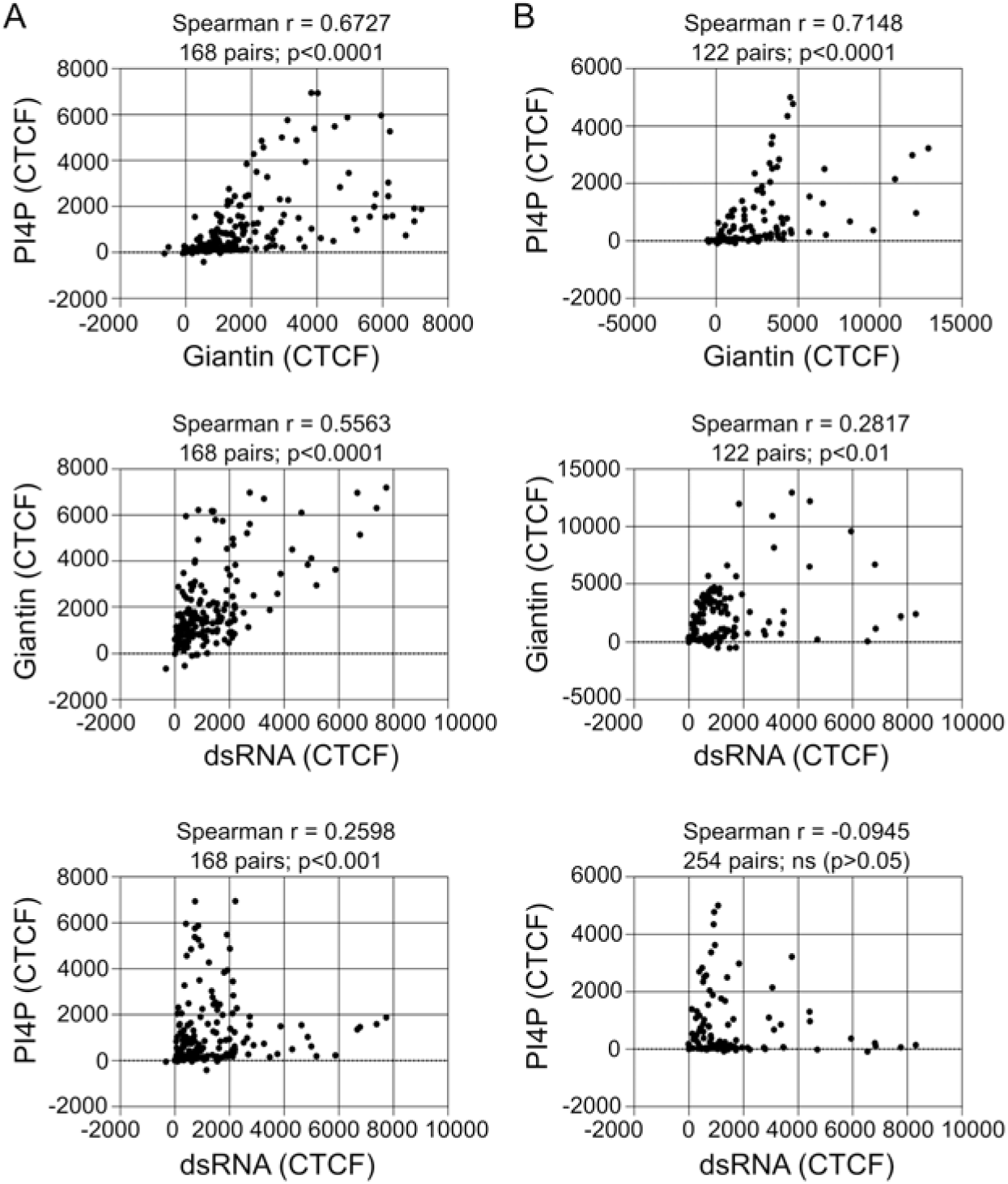
Neither the Golgi nor PI4P-positive vesicles are the main site for RV-C15 replication in HAE. **A-B:** Spearman correlation analysis (two-tailed; 0.95% confidence interval) between giantin and PI4P fluorescent levels (CTCF) in RV-A16 (**A**) or RV-A2 (**B**) -infected HAE at 12 hpi.

**S4 Figure.**
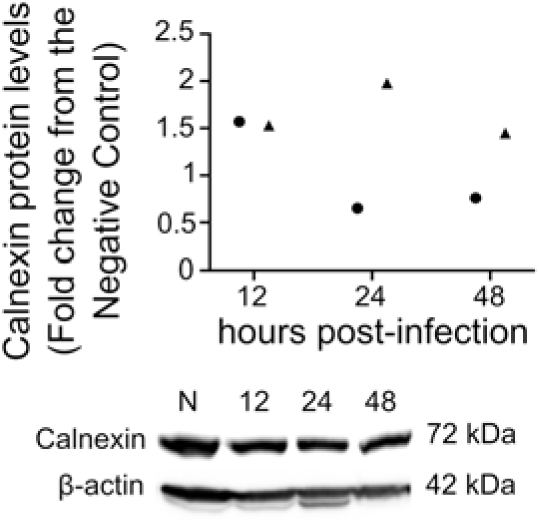
Global calnexin expression is not altered during RV-C15 infection in HAE. Fold change graph represents calnexin protein levels normalized to the endogenous control (actin) and compared to non-infected cultures. Data shown are from two independent donors, represented by circles and triangles. Blot below is from the donor represented by triangles.

**S5 Figure.**
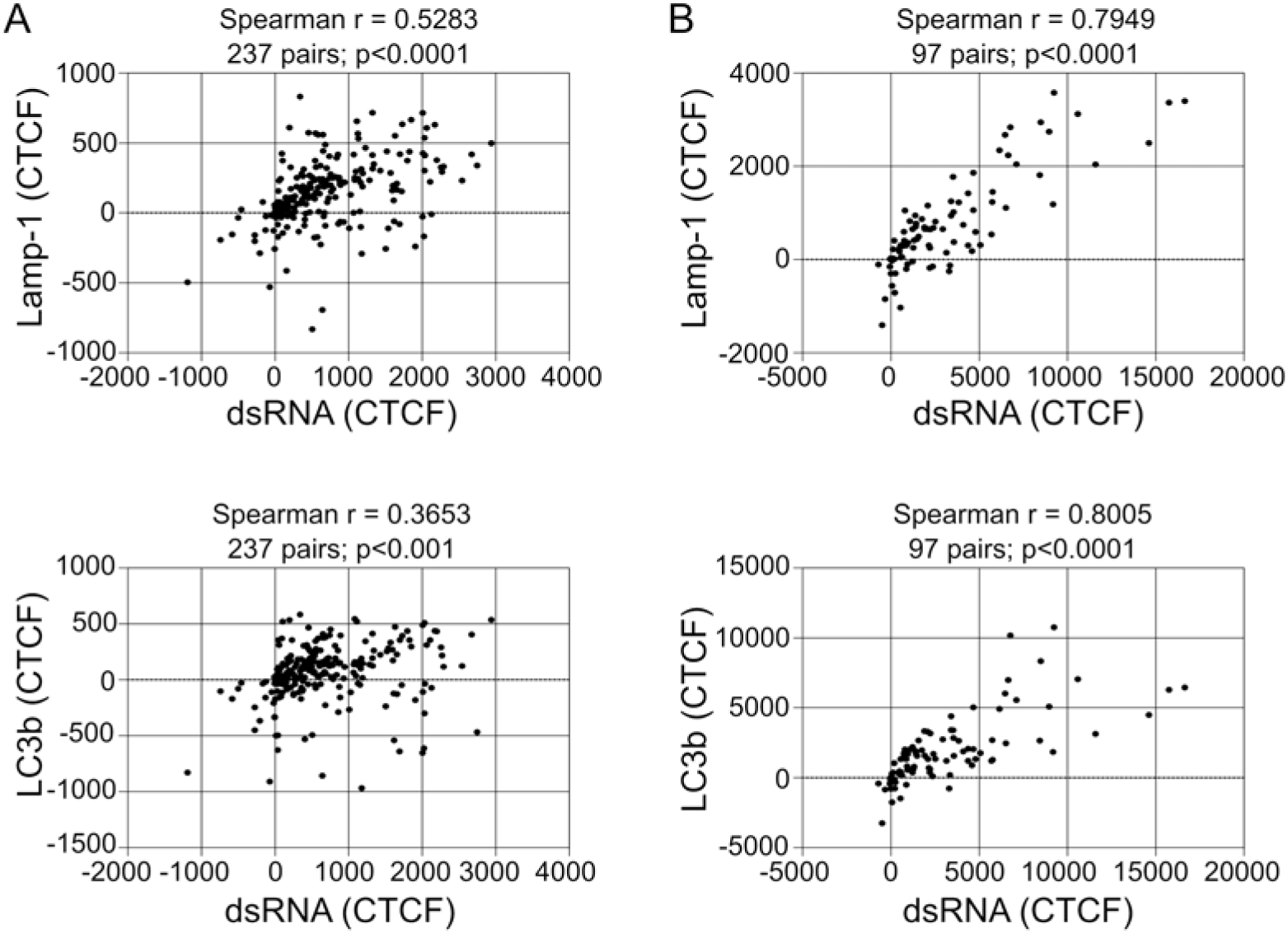
RV-C15 replication induces incomplete autophagy in HAE at 12 hpi. **A-B:** Spearman correlation analysis (two-tailed; 0.95% confidence interval) between dsRNA and Lamp-1 or LC3b fluorescent levels (CTCF) in RV-A16 (**A**) or RV-A2 (**B**) -infected HAE at 12 hpi.

**S6 Figure.**
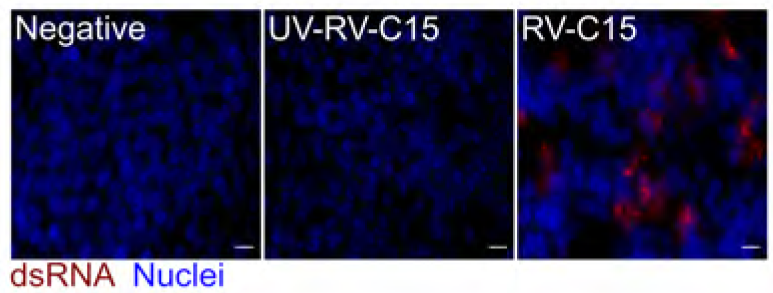
Detection of dsRNA in RV-C15, but not UV-RV-C15, inoculated HAE used in the MCC experiment. Immunofluorescence detection of dsRNA (red; nuclei, blue) in non-infected HAE or inoculated with UV- RV-C15 or RV-C15 at 12 hpi (scale bar = 10µm).

